# Early construction of the thalamocortical axon pathway requires JNK signaling within the ventral forebrain

**DOI:** 10.1101/2021.04.09.439208

**Authors:** Jessica G. Cunningham, James D. Scripter, Stephany A. Nti, Eric S. Tucker

## Abstract

Thalamocortical connectivity is essential for normal brain function. This important pathway is established during development, when thalamic axons extend a long distance through the forebrain before reaching the cerebral cortex. In this study, we identify a novel role for the c-Jun N-terminal Kinase (JNK) signaling pathway in guiding thalamocortical axons through intermediate target territories. Complete genetic removal of JNK signaling from the *Distal-less 5/6* (*Dlx5/6*) domain in mice prevents thalamocortical axons from crossing the diencephalon-telencephalon boundary (DTB) and the internal capsule fails to form. Ventral telencephalic cells critical for thalamocortical axon extension including corridor and guidepost neurons are also disrupted. In addition, corticothalamic, striatonigral, and nigrostriatal axons fail to cross the DTB. Analyses of different JNK mutants demonstrates that thalamocortical axon pathfinding has a non-autonomous requirement for JNK signaling. We conclude that JNK signaling within the *Dlx5/6* territory enables the construction of major axonal pathways in the developing forebrain.

**Impact statement:** We find an obligate, non-autonomous requirement for JNK signaling in early thalamocortical axon pathfinding, providing new mechanistic insight into the establishment of axonal connections in the developing forebrain.

## Introduction

The thalamocortical pathway is a major longitudinal axon projection that connects the diencephalon to the telencephalon and functions to relay essential sensory information to the cerebral cortex (Jones, 2001; Leyva-Diaz and Lopez-Bendito, 2013). A vital step in the formation of this connection occurs during embryonic development when growing thalamocortical axons cross the diencephalon-telencephalon boundary (DTB), an event assisted by chemorepulsion from the hypothalamus and chemoattraction to the ventral telencephalon (Lopez-Bendito and Molnar, 2003). The developing ventral telencephalon also serves as a critical intermediate target territory, containing corridor cells, guidepost neurons, and striatal axons that are necessary for proper guidance of thalamocortical axons (Lopez-Bendito et al., 2006; Molnar et al., 2012; Uemura et al., 2007). The correct patterning and cellular composition of this intermediate territory is critical for establishing proper thalamocortical connectivity, as disruptions to any one of these cellular populations or their axonal projections can result in the mistargeting of thalamocortical axons (Molnar et al., 2012).

The *Distal-less 5/6* (*Dlx5/6*) expression domain spans the entire axonal pathway through which thalamocortical axons project (Panganiban and Rubenstein, 2002), including the intermediate target territory of the ventral telencephalon. Interestingly, thalamocortical axons are misrouted in mice with conditional deletions of *Linx* (Mandai et al., 2014), *Ebf1* (Lokmane et al., 2013), *Celsr3* (Zhou et al., 2008; Zhou et al., 2009), or *Frizzled3* (Hua et al., 2014; Qu et al., 2014) from the *Dlx5/6* region. While these studies implicate the *Dlx5/6* territory as a critical intermediate target region for thalamocortical axons, downstream signaling pathways that could operate at the confluence of these different axon guidance systems are unknown.

The c-Jun N-terminal Kinase (JNK) signaling pathway has been implicated in axon development, including establishment of neuronal polarity, neurite architecture, axon guidance, and axon regeneration (Coffey et al., 2000; Oliva et al., 2006; Raivich et al., 2004; Schellino et al., 2019; Shafer et al., 2011; Tararuk et al., 2006). In mice, loss of *Jnk1* (Chang et al., 2003), the JNK scaffolding protein JIP3 (Cho et al., 2011; Kelkar et al., 2003), or upstream activators of JNK signaling including *DLK* (Hirai et al., 2006), *MKK4* (Wang et al., 2007), and *MKK7* (Yamasaki et al., 2011), lead to disruptions in the anterior commissure, corpus callosum, and internal capsule. The role for JNK in axon guidance within the *Dlx5/6* territory has not yet been explored, however our lab recently identified a novel role for JNK signaling in the migration of cortical interneurons arising from the *Dlx5/6* lineage (Myers et al., 2020; Myers et al., 2014; Smith et al., 2020). Since the *Dlx5/6* territory instructs thalamocortical axon pathfinding and disruptions to JNK signaling alter axonal development, we sought to determine whether JNK signaling was required within *Dlx5/6* cells to establish major longitudinal axon pathways in the forebrain.

Here, we demonstrate that complete loss of JNK function from the *Dlx5/6* territory of conditional triple knockout (*cTKO*) mice results in severe thalamocortical axon misrouting from early in development. In *cTKO* mice, thalamocortical axons misroute ventrally and rostrally instead of crossing into the ventral telencephalon at the DTB. Telencephalic corridor cells are mispositioned, and guidepost cells are reduced in number. Additionally, corticothalamic, striatonigral, and nigrostriatal axons are unable to project across the DTB and thus fail to reach their intended targets. This study is the first to demonstrate a non-autonomous requirement for JNK signaling in thalamocortical axon pathfinding and establishes critical roles for JNK signaling in axons and cell populations encompassed within the *Dlx5/6* territory.

## Results

### Axon tracts are missing or misrouted in JNK conditional triple knockout (cTKO) mice

Genetic disruption of the *Dlx5*/*6*-positive forebrain territory results in severe thalamocortical and corticothalamic axonal pathfinding defects (Hua et al., 2014; Mandai et al., 2014; Zhou et al., 2008). To determine whether JNK signaling is required for axon extension through this critical intermediate target territory, we utilized a conditional triple knockout (*cTKO*) mouse that conditionally removes *Jnk1* from *Dlx5/6-CRE-IRES-EGFP (Dlx5/6-CIE)* lineage cells of constitutive *Jnk2;Jnk3* double knockouts. Thus, JNK signaling is completely eliminated from the *Dlx5/6*-expressing territory through which major axon pathways traverse.

We began assessing axon tracts by labeling embryonic (E) day 17.5 brain sections with antibodies against L1, a transmembrane adhesion molecule expressed by many axons in the developing brain (Stallcup et al., 1985). In control brains at this age, the internal capsule contained L1-labeled axons connecting the cortex to the thalamus (Figure 1A,C, n=3/3). In *cTKO* mice however, the internal capsule was missing and no L1-labeled axons connected the diencephalon and telencephalon (Figure 1B,D, n=3/3). Instead, a large, “U-shaped” bundle of ectopically positioned axons was present beneath the hypothalamus (Figure 1B, n=3/3). Commissural axon tracts including the anterior commissure and corpus callosum were present in the *cTKO* forebrain and crossed the midline (Supplemental Figure 1A-D).

**Figure 1.**
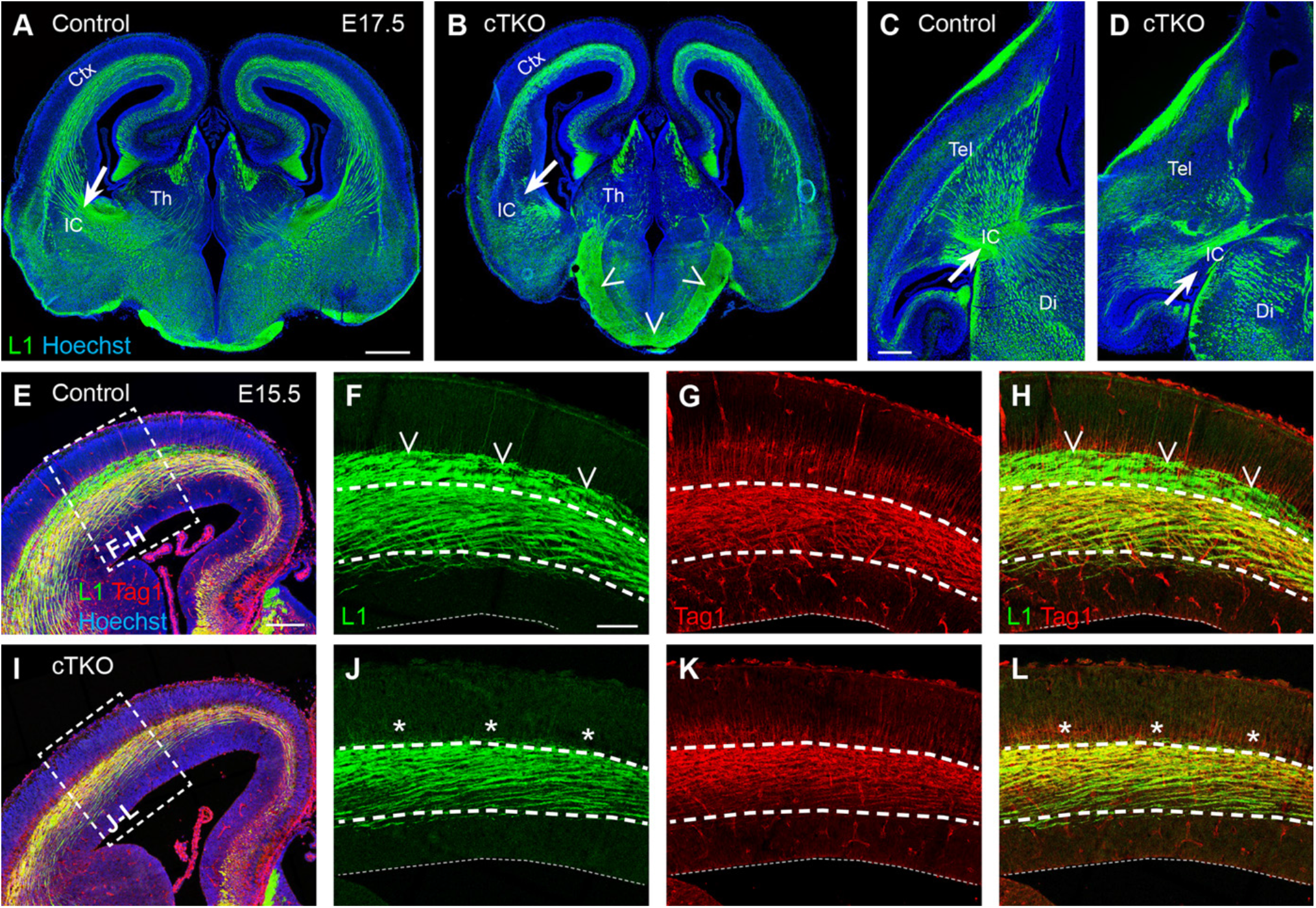
Axons are missing or misrouted in the *cTKO* brain. A-D. L1 labeling in coronal (A-B) and axial (C-D) sections of E17.5 brains. The internal capsule (arrow) is present in control (A,C) but not *cTKO* (B,D) brains. An ectopic “U-shaped” bundle projects into the hypothalamus of *cTKO* brains (open arrowheads, B). E-L. E15.5 cortices labeled with L1 and Tag1. L1-positve and Tag1-positive axons are present in control (E-H) and *cTKO* (I-L) cortices. A population of L1-positive/Tag1-negative axons is present in control brains (open arrowheads, F,H), but missing in *cTKO* brains (asterisks, J,L). Ctx-cortex, IC-internal capsule, Th-thalamus, Tel-telencephalon, Di-diencephalon. Scale bars: A-B 500 µm; C-D 300 µm; E,I 200 µm; F-H,J-L 100 µm.

*cTKO* cortices contained L1-labeled axons, however the abundance of these fibers appeared to be reduced. To determine the identity of these axons, we co-labeled E15.5 brain sections for L1 and Tag1, a neural adhesion molecule that marks corticofugal fibers (Wolfer et al., 1994) (Figure 1E-L). L1- and Tag1-positive axons were present in the intermediate zone (IZ) of both the control (Figure 1E-H; n=7/7) and *cTKO* (Figure 1I-L; n=7/7) cortical walls at E15.5. However, L1-positive/Tag1-negative fibers, which travel in the upper IZ, were completely absent from *cTKO* brains (Figure 1I-L, n=7/7), suggesting that a population of axons which originated sub-cortically was affected. Additionally, Calbindin-labeled axons were completely absent from *cTKO* brains at E15.5 (Supplemental Figure 1E-F, n=11/11). Thus, we found several major axon deficiencies in *cTKO* forebrains, including a disrupted internal capsule, an ectopic accumulation of axonal fibers coursing beneath the hypothalamus, and a loss of axons normally traversing the upper IZ in the cortical wall.

### Thalamocortical axons misroute ventrally in cTKO mice

Since L1-postive/Tag-1 negative axons were absent from *cTKO* cortices and there was an ectopic bundle of L1-positive axons beneath the hypothalamus, we hypothesized that thalamocortical axons were misrouted in *cTKO* brains. A DiI crystal was placed into the dorsal thalamus of fixed brain slices at E15.5 to observe the projection pattern of thalamocortical axons (Figure 2A). In control slices, DiI-labeled axons extended from the thalamus, crossed the diencephalon-telencephalon boundary (DTB), and projected through the internal capsule in the telencephalon (Figure 2B, n=8/8). However, in *cTKO* slices, DiI-labeled axons projected to the DTB, but rather than crossing into the telencephalon at this level, turned ventrally and coursed through the hypothalamus (Figure 2C, n=3/3).

**Figure 2.**
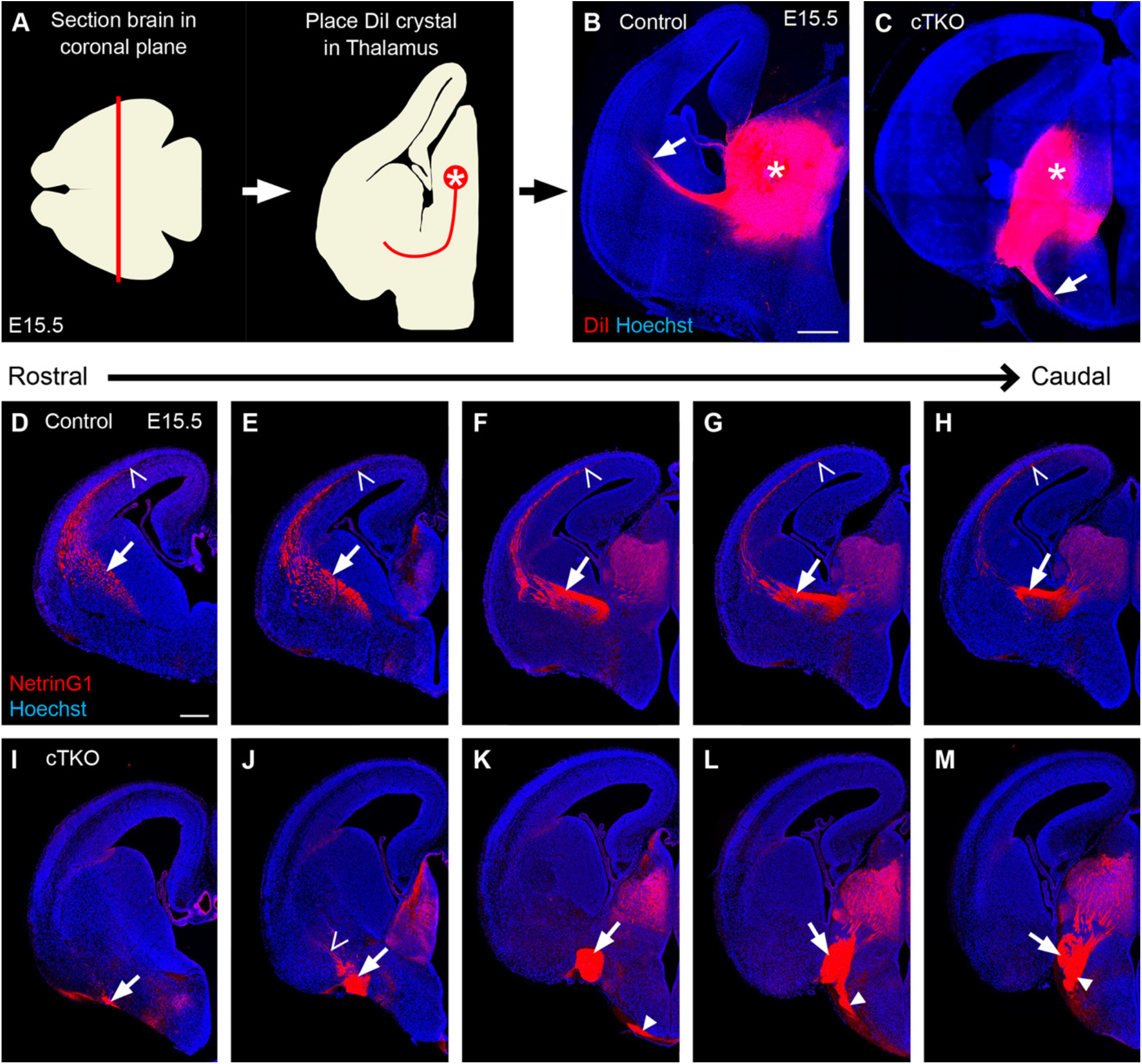
Thalamocortical axons are misrouted in *cTKO* brains. A. E15.5 brains were fixed, sectioned coronally at 300 microns, and a DiI crystal (asterisk) was placed into the dorsal thalamus. B-C. DiI labels axons projecting through the internal capsule of control brains (arrow, B), and axons projecting ventrally into the hypothalamus of *cTKO* brains (arrow, C). D-M. NetrinG1-labeled thalamocortical axons in rostrocaudal series of control (D-H) and *cTKO* (I-M) brains at E15.5. In control brains, axons project through the internal capsule (arrow, D-H) and reach the cortex at all levels (open arrowheads, D-H). In *cTKO* brains, fibers aberrantly project ventrally into the hypothalamus (closed arrowheads, K-M), rostrally along the ventromedial margin of the telencephalon (arrows, I-M), and into the striatum at ectopic locations (open arrowhead, J). Scale bars: 300 µm.

To further assess the trajectory of thalamocortical axons, we labeled E15.5 brain sections for NetrinG1, a netrin that is enriched in the dorsal thalamus during development and serves as a marker of most thalamocortical axons (Nakashiba et al., 2000). In control brains, NetrinG1-positive thalamocortical axons crossed the DTB at caudal levels, extended through the internal capsule, and reached the mid-cortical wall at all rostrocaudal levels examined (Figure 2D-H, n=4/4). However, in *cTKO* brains, thalamocortical axons extended from the thalamus and projected ventrally into the hypothalamus instead of crossing the DTB (Figure 2K-M, n=15/15). Additionally, a second bundle of misrouted thalamocortical axons extended rostrally in a large fascicle located at the ventromedial margin of the telencephalon in *cTKO* brains (Figure 2I-M, n=15/15). From this rostrally-projecting bundle, some thalamocortical axons splayed into the ventral forebrain at ectopic positions (Figure 2J), but with varying degrees of penetration (Supplemental Figure 2). Together, analyses of both DiI and NetrinG1 labeling indicate that thalamocortical axons are massively misrouted at the DTB region in developing *cTKO* brains.

#### Thalamocortical and corticothalamic axons take aberrant trajectories in cTKO mice

Immunohistochemical analysis revealed that thalamocortical axons mis-projected both ventrally and rostrally in *cTKO* brains at E15.5. Despite failing to cross the DTB and properly extending through the internal capsule region, some NetrinG1-labeled thalamic projections entered the telencephalon at aberrant locations (Supplemental Figure 2, n=9/15 NetrinG1-labeled brains). To determine whether these fibers eventually extended into the cortex, we examined brains at postnatal (P) day 0, the oldest age we were able to collect viable *cTKO* mice (Table 1). In addition, we asked whether the subcortical trajectory of cortical axons was also compromised in *cTKO* mice, since thalamocortical and corticothalamic axons are known to be intimately associated with one another during development (Chen et al., 2012; Molnar et al., 1998). P0 brains were collected, fixed, hemisected along the midsagittal plane, and a DiI crystal was placed into the thalamus of one hemisphere (Figure 3A) and into the cortex of the opposite hemisphere (Figure 4A) of each brain.

**Table 1.** Offspring from experimental cross used to generate *cTKO* mice.

| Age: | Litters: | Total Animals: | *cTKO*: | Percent: |
| --- | --- | --- | --- | --- |
| E12.5 | 4 | 32 | 5 | 15.6% (5/32) |
| E13.5 | 19 | 149 | 19 | 12.8% (19/149) |
| E14.5 | 1 | 8 | 1 | 12.5% (1/8) |
| E15.5 | 31 | 208 | 28 | 13.5% (28/208) |
| E17.5 | 8 | 52 | 10 | 19.2% (10/52) |
| P0 | 33 | 216 | 30 | 13.9% (30/216) |
| P21 | 10 | 43 | 0 | 0.0% (0/43) |

Numbers of litters, total animals, and *cTKO* mice recovered between embryonic (E) day 12.5 and postnatal (P) day 21. Expected ratios (1/8) of *cTKO* mice were collected at all time points except P21.

**Figure 3.**
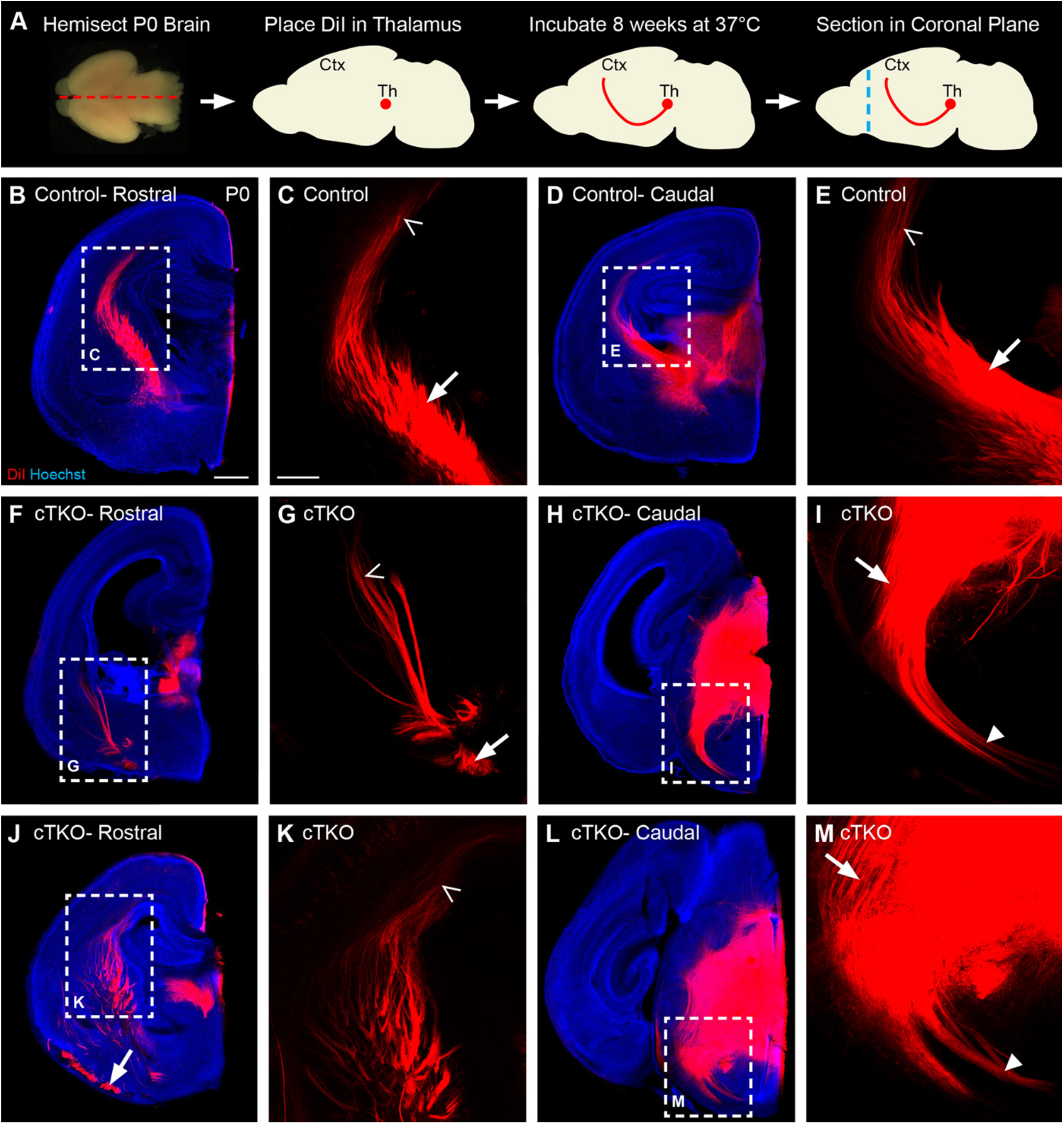
Axons projecting from the thalamus have aberrant trajectories in the P0 *cTKO* brain. A. P0 brains were hemisected, DiI-labeled in the thalamus, incubated, and coronally sectioned at 200 microns. B-E. DiI-labeled axons in control brains project through the internal capsule (arrow) to reach the cortex (open arrowhead) at rostral (B, C) and caudal (D, E) locations. F-M. DiI-labeled axons in *cTKO* brains project ventrally at caudal locations (arrow, I,M) and extend into the hypothalamus (closed arrowhead, I,M). At more rostral levels (F-G, J-K), a bundle of axons resides along the ventral margin of the telencephalon (arrow, G,J). DiI-labeled axons in some *cTKO* brains stop in the striatum (open arrowhead, G), while DiI-labeled axons in other *cTKO* brains reach the cortex (open arrowhead, K) in reduced numbers. Scale bars: B,D,F,H,J,L 500 µm; C,E,G,I,K,M 200 µm.

**Figure 4.**
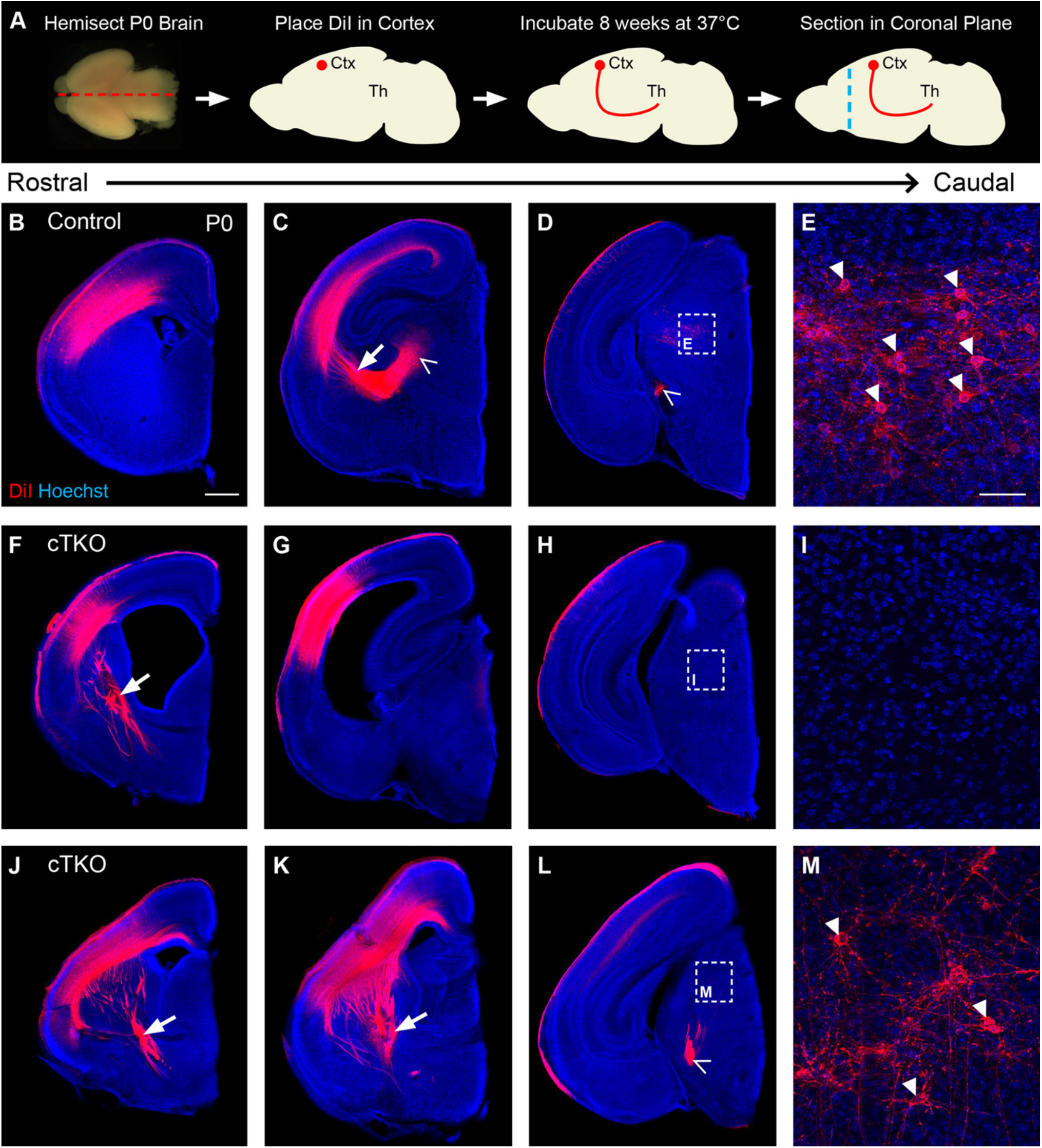
Corticothalamic axons display aberrant projection patterns in the P0 *cTKO* brain. A. P0 brains were hemisected, DiI-labeled in the cortex, incubated, and coronally sectioned at 200 microns. B-E. DiI-labeled cortical axons in control brains project through the internal capsule (arrow, C) and into the thalamus (open arrowheads, C-D). Cell bodies of retrogradely labeled thalamocortical neurons are present in the thalamus (closed arrowheads, E). F-M. DiI-labeled cortical axons in *cTKO* brains project rostrally and ventrally in large torturous fascicles (arrows, F,J-K). In n=2/4 *cTKO* brains (F-I), no cortical axons reach the thalamus (G-H), and no retrogradely labeled cells are present (I). In n=2/4 *cTKO* brains (J-M), axons reach the thalamus (open arrowhead, L), and retrogradely labeled cells are present (closed arrowheads, M). Scale bars: B-D, F-H, J-L 500 µm; E,I,M 50 µm.

In control brains, DiI-labeled axons projected from the thalamus, across the DTB, through the internal capsule, and into the cortex at rostral and caudal locations (Figure 3B-E; n=5/5). However, in all *cTKO* brains examined, thalamocortical axons misrouted ventrally into the hypothalamus (Figure 3H-I, L-M, n=4/4) and rostrally along the ventromedial margin of the forebrain (Figure 3F-G, J-K). This finding verified our NetrinG1 (Figure 2M) and DiI-labeled slice (Figure 2D) results at E15.5. Similar to the E15.5 timepoint, thalamocortical axons were observed invading the striatum from an ectopic ventral position at P0 in *cTKO* brains (Figure 3F-G, J-K). These ectopically projecting DiI-labeled fibers only reached the cortex in half of the *cTKO* brains examined (Figure 3J, n=2/4), and when they did, they were less abundant and less advanced than their control counterparts. We also noticed increased ventricular volume in *cTKO* slices, which has been reported in other mouse mutants with thalamocortical axon guidance phenotypes (Hua et al., 2015; Hua et al., 2014; Tissir et al., 2005; Wang et al., 2002). CT scans were performed on whole heads at P0, where we observed increased ventricular volume in the intact *cTKO* brain (Supplemental Figure 3, n=6/6).

We next examined corticothalamic projections in control and *cTKO* brains (Figure 4). In control brains, DiI-labeled corticothalamic axons assumed a normal trajectory through the internal capsule, crossed into the diencephalon, and reached the dorsal thalamus (Figure 4B-D, n=5/5). Additionally, DiI-labeled cell bodies were present in the thalamus from retrograde labeling of thalamocortical axons (Figure 4D-E, n=5/5). In *cTKO* brains however, corticofugal axons were severely misrouted and did not coalesce into an organized internal capsule (Figure 4F-G,J-K, n=4/4). Instead, cortical axons ectopically entered the subpallium in large, disorganized fascicles at rostral levels (Figure 4F,J-K, n=4/4). Cortical fibers only reached the thalamus in half of the *cTKO* brains examined (Figure 4L, n=2/4). Interestingly, retrogradely labeled thalamic cells were only found in *cTKO* brains that contained corticothalamic fibers in the thalamus, and when present, were reduced in number (Figure 4L-M).

Since a DiI crystal was placed into either the thalamus (Figure 3) or cortex (Figure 4) of the same hemisected brain, we compared the degree of axon misrouting between the thalamic and cortical DiI placements. Indeed, we found that in the *cTKO* brains that lacked thalamic innervation of the cortex (Figure 3F-I), no cortical axons had reached the thalamus (Figure 4F-G). Additionally, no retrogradely labeled thalamic cells were present after the cortical DiI placement (Figure 4H-I), indicating that thalamic axons had not reached the cortex in either hemisphere of each brain examined. Similarly, in *cTKO* brains with partial thalamic innervation of the cortex (Figure 3J-M), some cortical axons had reached the thalamus (Figure 4J-L), and retrogradely labeled thalamic cells were present (Figure 4L-M). Thus, reciprocal thalamocortical and corticothalamic projections were comparably affected in each hemisphere of the *cTKO* brains examined.

#### Thalamocortical axons are misrouted early in development in cTKO brains

We next assessed the developmental time course of thalamocortical axon misrouting in *cTKO* brains. Using NetrinG1 as a marker of thalamocortical axons, we examined brains ranging from E12.5 to P0 (Figure 5A-J). At E12.5, when thalamocortical axons first cross from the diencephalon into the telencephalon (Figure 5A inset, n=2/2), no NetrinG1-positive fibers crossed the DTB in *cTKO* mice (Figure 5F inset; n=2/2). Similarly, at E13.5 when the furthest advanced thalamocortical fibers have approached the pallial-subpallial boundary in control brains (Figure 5B, n=6/6), NetrinG1-positive axons were absent from the ventral telencephalon in *cTKO* mice (Figure 5G, n=8/8). When we examined the diencephalon-telencephalon boundary (DTB) region at E13.5, where NetrinG1-positive axons normally cross the DTB in control brains (Figure 5K-L, n=6/6), we observed ventrally projecting thalamocortical axons in *cTKO* brains that failed to cross the DTB (Figure 5M-N, n=8/8). Thus, the misrouting of thalamocortical axons begins in early development, when thalamocortical axons would normally first cross the DTB region. Once misrouted, thalamocortical axons remain similarly positioned in *cTKO* brains at E15.5 (Figure 5H, n=15/15), E17.5 (Figure 5I, n=3/3), and P0 (Figure 5J, n=5/5), with the size of the “U-shaped” bundle increasing in thickness as development proceeds.

**Figure 5.**
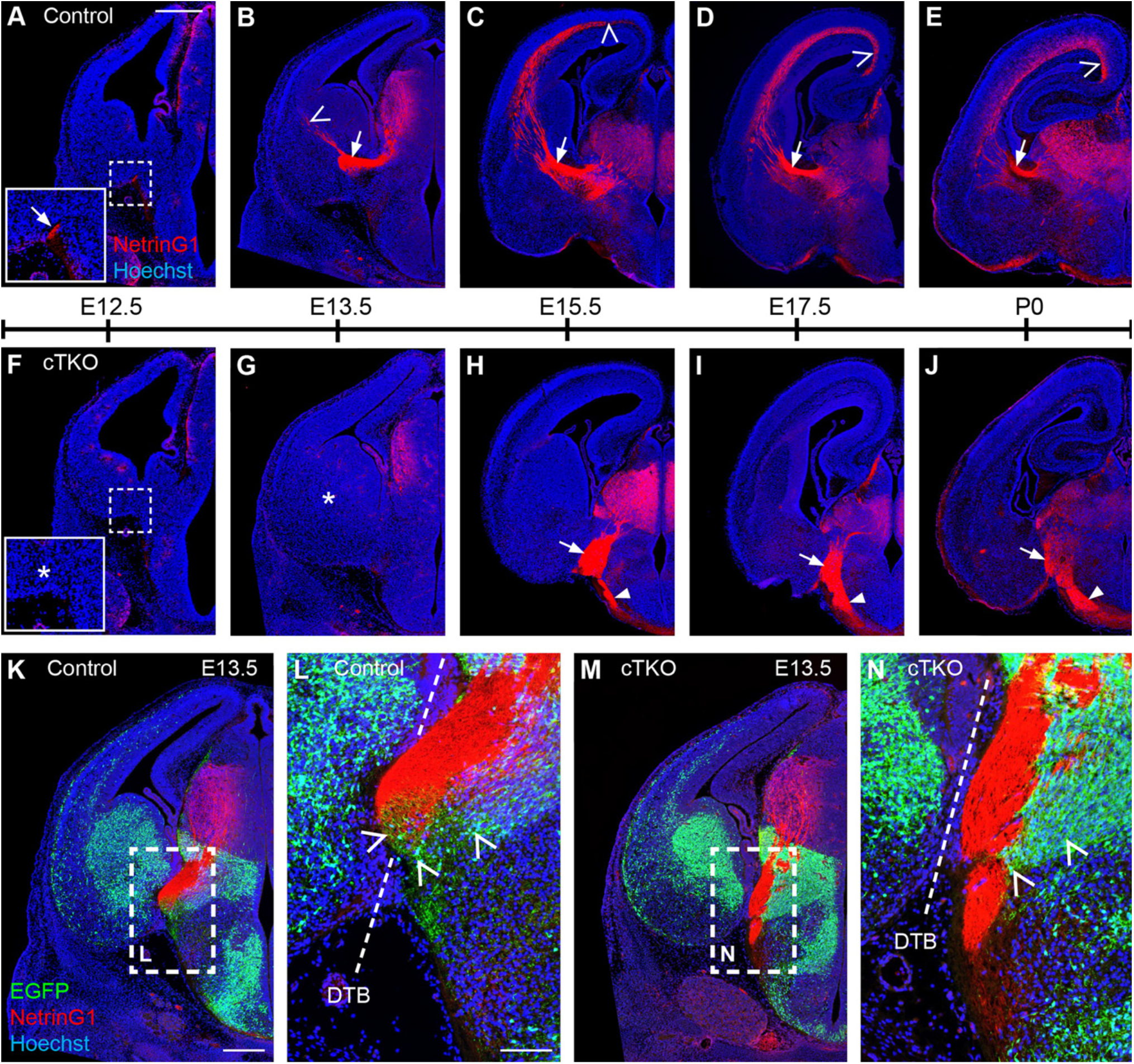
Thalamocortical axons are misrouted from early developmental timepoints in the *cTKO* brain. A-E. NetrinG1 labeling of thalamocortical axons from E12.5 to P0 in control brains at the level of the diencephalon-telencephalon boundary (DTB). Axons extend into the ventral telencephalon at E12.5 (arrow-inset, A), reach the pallial-subpallial boundary at E13.5 (open arrowhead, B), and continue to extend through the internal capsule (arrows, B-E) and into the cortex from E15.5-P0 (open arrowheads, C-E). F-G. In *cTKO* brains, NetrinG1-positive axons are missing in the ventral telencephalon at E12.5 (asterisk, F) and E13.5 (asterisk, G). H-J. Axons are present adjacent to the DTB (arrows) or below the hypothalamus (closed arrowheads) in *cTKO* mice from E15.5-P0. K-L. NetrinG1-positive thalamocortical axons project from the thalamus and cross into the ventral telencephalon at the DTB (dashed line, L) in close association with *Dlx5/6*-positive cells and axons (open arrowheads, L) at E13.5. M-N. In *cTKO* brains, thalamocortical axons have extended from the thalamus, however they have not crossed the DTB (dashed line, N). *Dlx5/6-CIE* cells and axons fail to span the DTB and remain positioned in the diencephalon (open arrowheads, N). Scale bars: scale bar in A represents 400 µm for A,B,F,G, 500 µm for C,D,H,I, and 600 µm for E,J; K,M 300 µm, 100 µm L,N.

In E13.5 *cTKO* brains, thalamocortical axons extended ventrally from the thalamus and through the prethalamus (Figure 5M-N, n=8/8), suggesting their initial extension in the diencephalon is grossly unaffected. However, after exiting the prethalamus, thalamocortical axons were unable to cross the DTB. Interestingly, this pathway is populated by *Dlx5/6-CIE*-positive cells and axons, which form a bridge from the prethalamus to the ventral forebrain in control brains (Figure 5K-L, n=6/6). In *cTKO* brains however, *Dlx5/6-CIE*-positive cells were present either in the telencephalon or diencephalon, but like the thalamocortical axons, they failed to span the DTB (Figure 5M-N, n=8/8). Indeed, *Dlx5/6-CIE*-positive cells appeared to form a piled-up wedge on the diencephalic side of the DTB in *cTKO* brains (Figure 5M-N, n=8/8). These data collectively suggest that thalamocortical axons are able to extend properly from the thalamus to the DTB in *cTKO* brains, however at the DTB region, there is a disruption to both thalamocortical axons and the *Dlx5/6-CIE*-positive cells and axons that normally bridge this territory.

#### Corridor and guidepost cells are disrupted in cTKO mice

The corridor is a permissive territory in the developing ventral telencephalon that supports the extension of thalamocortical axons through an otherwise growth inhibitory environment (Bielle et al., 2011; Lopez-Bendito et al., 2006). Since disruptions to the corridor region have been implicated in thalamocortical axon misrouting (Molnar et al., 2012), and the *Dlx5/6* domain spans this region, we examined the corridor in control and *cTKO* brains at E13.5 at 4 different rostro-caudal locations (Figure 6). Brain slices were labeled for Islet1 to reveal corridor cells (Figure 6B-E), which migrate ventral-medially from the lateral ganglionic eminence (LGE) to position themselves between the Nkx2.1-positive medial ganglionic eminence (MGE) progenitor domain and the nascent globus pallidus (GP). The Islet1-positive corridor also extends caudally to the DTB, where thalamocortical axons project from the diencephalon into the telencephalon (Figure 6D-E, n=4/4). In *cTKO* brains, the distribution of Islet1-positive cells was similar to control brains at rostral levels (Figure 6F, n=5/5). However, immediately caudal to this location, Islet1-positive cells were ectopically positioned laterally and ventrally, in the region where the GP is typically forming (Figure 6G, n=5/5). At more caudal levels, where Islet1-positive cells normally associate with thalamocortical axons in control brains (Figure 6D-E, n=4/4), the Islet1-positive cells were severely reduced in number in *cTKO* brains in the ventral telencephalon, and no thalamocortical axons had crossed the DTB (Figure 6H-I, n=5/5).

**Figure 6.**
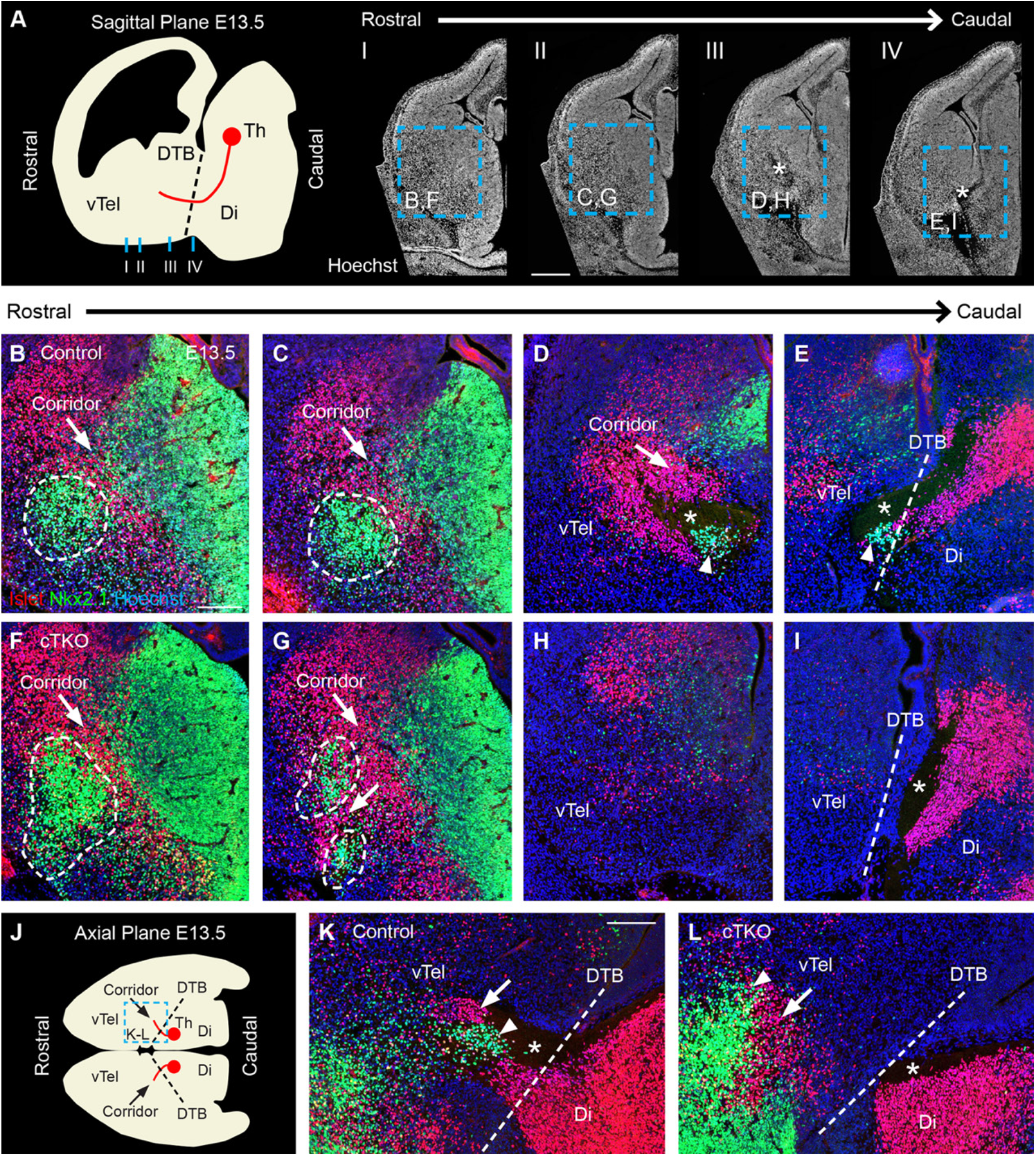
The ventral telencephalic corridor is malformed in *cTKO* brains at E13.5. A. Schematic and Hoechst-labeled slices illustrating positions used for Islet1 and Nkx2.1 analysis. B-D. Islet1-positive (arrows) cells form a permissive corridor between the Nkx2.1-positive medial ganglionic eminence and globus pallidus (encircled B-C) through which unlabeled thalamocortical axons extend (asterisk, D). E. Thalamocortical axons (asterisk) cross the DTB (dashed line) above a patch of Nkx2.1-positive cells (closed arrowhead) and Islet1-positive cells from the diencephalon. F-I. In *cTKO* brains, the Islet1-positive corridor is present at rostral levels (arrows, F-G), but reduced at caudal levels (H-I). The Nkx2.1-positive globus pallidus is expanded rostrally (encircled, F), ectopically infiltrated by Islet1-positive cells at mid levels (arrow, G), and diminished caudally (H-I) in *cTKO* brains. I. Thalamocortical axons (asterisk) fail to cross the DTB. Islet1-positive cells are present in the diencephalon, but are missing at the DTB (dashed line). J-L. Axial view of the E13.5 corridor. Islet1*-* (arrows) and Nkx2.1-(closed arrowheads) positive cells extend caudally to where thalamocortical axons (asterisk) cross the DTB (dashed line) in controls (K), but stop rostrally in *cTKO* brains (L). vTel-ventral telencephalon, Di-diencephalon, DTB-diencephalon-telencephalon boundary. Scale bars: A 400 µm; B-I, K-L 150 µm.

Islet1-positive cells located in the telencephalon as well as the diencephalon are critical for laying down a pioneering scaffold which acts as a bridge across the DTB for early thalamocortical axons (Feng et al., 2016). The diencephalic Islet1-positve cells in the prethalamus are *Dlx5/6*-positive, and fail to extend across the DTB in mutants where thalamic axons also fail to cross this boundary (Feng et al., 2016). In control brains, diencephalic Islet1-positive cells formed a scaffold beneath the thalamocortical axons (Figure 6E,K, n=4/4). Unlike telencephalic Islet1-positive cells that were depleted at this level in *cTKO* brains, diencephalic Islet1-positive cells were present in seemingly normal numbers in *cTKO* mice (Figure 6I,L, n=5/5). However, these Islet1-positive cells stopped short of the DTB at the position where thalamocortical axons misrouted ventrally (Figure 6I,L, n=5/5).

We next examined the Nkx2.1 expression domain, which is altered in other knockout mice with thalamocortical axon misrouting phenotypes (Jia et al., 2014; Lakhina et al., 2007; Tuttle et al., 1999; Uemura et al., 2007). In control brains, Nkx2.1-positive GP cells were present throughout the corridor region and resided directly beneath the thalamocortical axons as they entered the telencephalon (Figure 6D-E, n=4/4). However, in *cTKO* brains, Nkx2.1-positive GP cells occupied a larger territory at rostral levels, extending to the lateral, pial margin of the ventral telencephalon (Figure 6F, n=5/5). Slightly more caudal, the Nkx2.1-positive GP domain shifted dorsal and lateral, in some cases splitting into multiple clusters with the apparent influx of ectopically positioned Islet1-positive cells (Figure 6G, n=5/5). Similar to Islet1-positive cells, Nkx2.1-positive cells were missing from the caudal corridor of *cTKO* brains, where thalamocortical axons normally cross into the telencephalon (Figure 6H-I, n=5/5). In control brains, both Islet1- and Nkx2.1-positive cells were precisely positioned at the point of axon crossing at the DTB (Figure 6E,K, n=4/4), however, these two populations of cells failed to extend caudally to the DTB in *cTKO* brains (Figure 6I,L, n=5/5), which can be observed in both coronal and axial planes.

Another population of cells thought to assist thalamocortical axons across the DTB are guidepost cells (Braisted et al., 1999), which reside in the ventral telencephalon and extend axons to the dorsal thalamus. Since these cells are often missing or reduced in number in mutants with thalamocortical axon defects (Molnar et al., 2012), and are located within the *Dlx5/6* territory, we investigated whether they were present in the brains of *cTKO* mice. While the molecular profile of these guidepost cells remains unknown, they can be retrogradely labeled by placing a DiI crystal into the dorsal thalamus (Figure 7). In E13.5 control brains, thalamocortical axons crossed the DTB and projected through the ventral telencephalon (Figure 7A-D; n=6/6). Retrogradely labeled cells were found in close association with the internal capsule at both rostral (Figure 7A-B) and caudal (Figure 7C-D) levels of the ventral telencephalon in territories that were *Dlx5/6-CIE* positive. In *cTKO* brains, thalamocortical axons projected ventrally from the thalamus, but did not cross the DTB and failed to reach the telencephalon (Figure 7E-H, n=6/6). Retrogradely labeled cells were significantly reduced in number in *cTKO* brains at both rostral (Figure 7E-F, Control: 46.0 ± 5.0, *cTKO*: 12.3 ± 4.5, Student’s T-Test p=0.0005, n=6/6) and caudal (Figure 7G-H, Control: 20.0 ± 4.6, *cTKO*: 1.83 ± 0.70, Student’s T-Test p=0.0029, n=6/6) locations. Collectively, both corridor and guidepost neurons, which are known to play a critical role in early thalamocortical axon pathfinding and are present in the *Dlx5/6* domain, are disrupted in *cTKO* brains.

**Figure 7.**
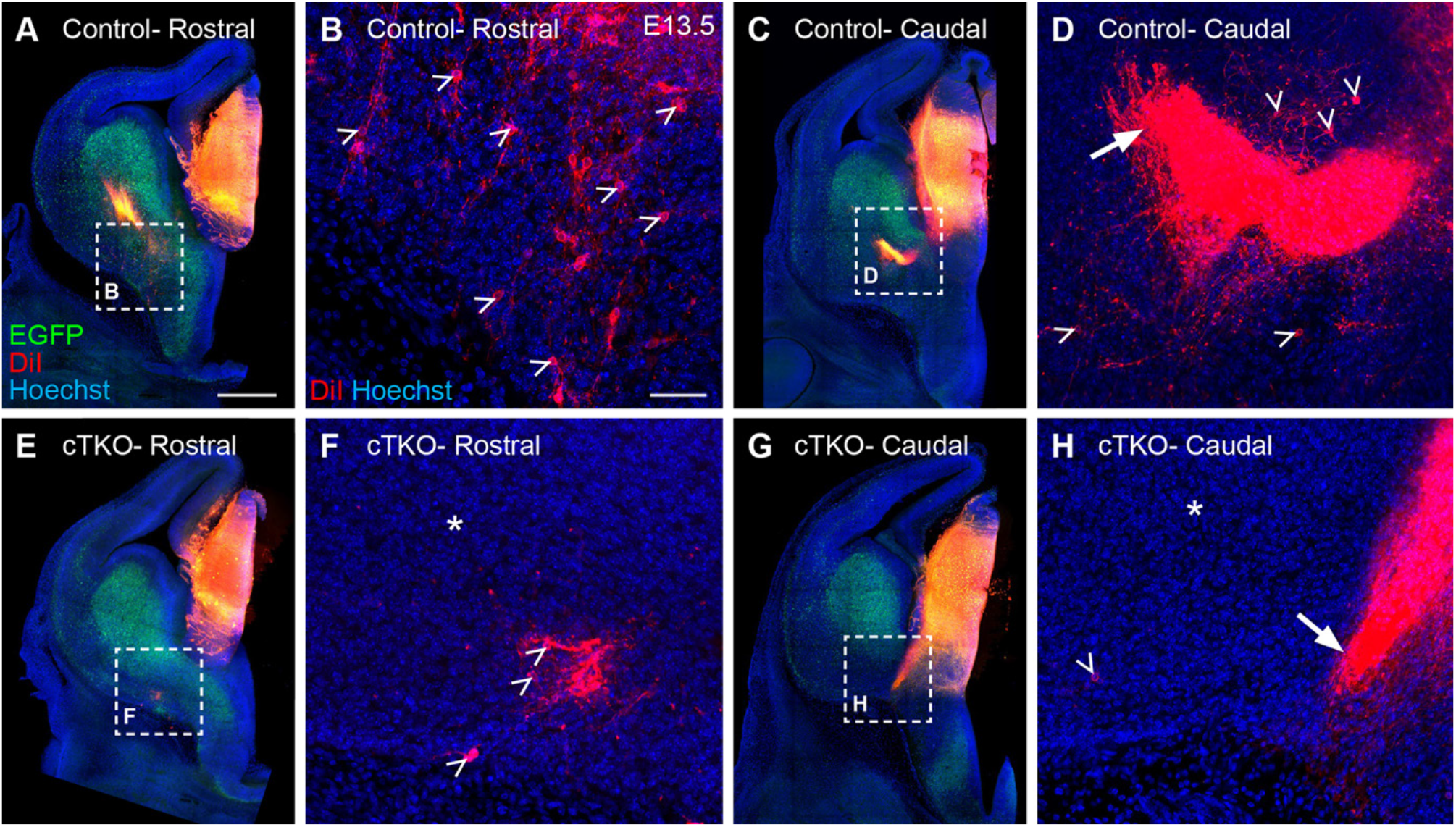
Ventral telencephalic guidepost cells are reduced in number in *cTKO* brains at E13.5. A-D. DiI-labeled thalamocortical axons (arrow, D) and retrogradely-labeled guidepost cells (open arrowheads, B,D) are present in the telencephalon after DiI placement in the dorsal thalamus at E13.5 in control brains. E-H. In *cTKO* brains, thalamocortical axons are missing in the telencephalon (asterisks, F,H), and instead project ventrally (arrow, H). A few retrogradely-labeled guidepost cells are present in the *cTKO* ventral telencephalon, but are reduced in number (open arrowheads, F,H). Scale bars: A,C,E,G 400 µm, B,D,F,H 50 µm.

### Reciprocal connections between the striatum and substantia nigra are altered in cTKO mice

Besides thalamocortical and corticothalamic axons, several other axonal pathways cross the DTB during early development. Two pathways in particular reciprocally connect the striatum to the substantia nigra (striatonigral) and the substantia nigra to the striatum (nigrostriatal). Since these pathways course in close proximity to thalamocortical axons as they approach the DTB, and disruptions to both the striatonigral (Jia et al., 2014; Morello et al., 2015; Uemura et al., 2007) and nigrostriatal (Bagri et al., 2002; Fenstermaker et al., 2010; Uemura et al., 2007) pathways have been observed in mice with thalamocortical axon guidance defects, we examined striatal and nigral axon projections in *cTKO* mice. In control brains, Darpp32-positive striatal axons projected from the striatum, crossed the DTB, and extended to the substantia nigra in the midbrain (Figure 8A,C-E,I n=6/6). Reciprocally, tyrosine hydroxylase (TH)-positive nigral axons projected from the substantia nigra to join the median forebrain bundle, crossed the DTB, and distributed diffusely in the striatum (Figure 8A,I n=6/6). However, in *cTKO* brains, Darpp32-positive fibers coalesced and became entangled in the striatum (Figure 8B,F; n=5/5). These striatal fibers did not cross the DTB and were therefore missing at the level of the diencephalon and substantia nigra (Figure 8B,G-H,J). TH-positive nigrostriatal fibers were likewise unable to fully extend along the anterior-posterior axis in *cTKO* brains (Figure 8B), and instead mainly misrouted in a “U-shaped” bundle into the hypothalamus (Figure 8J, n=8/8), similar to the NetrinG1-positive thalamocortical axons. In both the thalamocortical and nigrostriatal pathways, we observed similar patterns of misrouting, in which normally ipsilaterally projecting tracts were misrouted across the ventral midline. We next wanted to determine if nigrostriatal axons mis-projected early in development similar to thalamocortical axons. Indeed, TH-positive axons in *cTKO* mice were misrouted as early as E13.5 (Supplemental Figure 4F, n=4/4), and had formed a similar “U-shaped” bundle by E15.5 (Supplemental Figure 4L, n=15/15). In *cTKO* mice, multiple longitudinal axon pathways are unable to properly extend ipsilaterally along the rostrocaudal axis, fail to cross the DTB, and at least some of these fibers incorrectly project across the midline.

**Figure 8.**
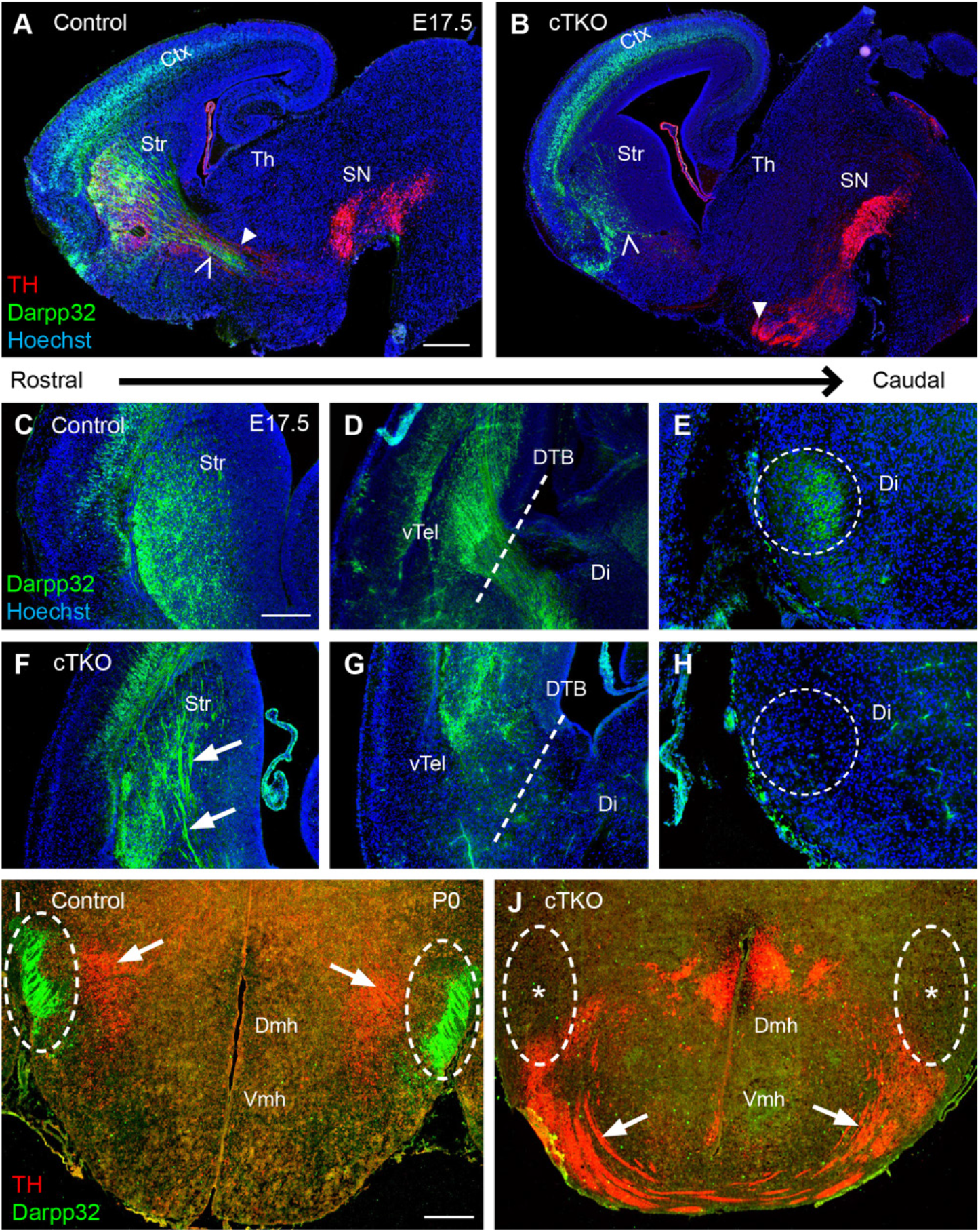
Axons fail to interconnect the striatum and the substantia nigra in *cTKO* brains. A. In sagittal sections of E17.5 control brains, Darpp32-positive striatonigral axons (open arrowhead) and TH-positive nigrostriatal axons (closed arrowhead) project across the DTB in control brains. B. In E17.5 *cTKO* brains, Darpp32-postive axons (open arrowhead) fail to cross the DTB and TH-positive axons (closed arrowhead) misproject into the hypothalamus. C-E. In coronal sections of E17.5 control brains, Darrp32-positve axons are present in the striatum (C), cross the DTB (D), and reach the diencephalon (E). F-H. In E17.5 *cTKO* brains, Darrp32-positive axons are misrouted in the striatum (arrows, F), do not cross the DTB (G), and are absent in the diencephalon (H). I. In coronal sections of P0 control brains, Darpp32 (encircled) and TH axons (arrows) travel adjacently in the hypothalamus. J. In P0 *cTKO* rains, Darpp32-positive axons are missing (asterisks), and TH-positive axons misroute ventrally (arrows). Ctx- cortex, Str- striatum, Th- thalamus, SN- substantia nigra, vTel- ventral telencephalon, Di-diencephalon, DTB- diencephalon-telencephalon boundary, Dmh- dorsomedial hypothalamic nucleus, Vmh- ventromedial hypothalamic nucleus. Scale bars: A-B, 400 µm; C-J, 250µm.

### Non-autonomous requirement for JNK signaling in thalamocortical axon pathfinding

Although our data strongly implicate a non-autonomous role for JNK signaling in establishing major forebrain axon tracts, we cannot exclude the possibility that the constitutive loss of *Jnk2* and *Jnk3* in *cTKO* mice impacts thalamocortical projections. In *cTKO* mice, *Jnk1* is conditionally removed from *Dlx5/6-CIE* cells in a *Jnk2;Jnk3* double knockout background (Figure 9B), where thalamocortical axons themselves have lost *Jnk2* and *Jnk3* function. To test whether the loss of *Jnk2* and *Jnk3* alone impairs thalamocortical axon pathfinding, we developed a new mouse line to constitutively remove *Jnk2* and *Jnk3*, while leaving *Jnk1* function intact (Figure 9C). NetrinG1-positive thalamocortical axons in *Dlx5/6-CIE;Jnk2-/-; Jnk3-/-* mice extended through the internal capsule and reached the cortex at E15.5 (Figure 9C, n=3/3) and at P0 (Supplemental Figure 5C, n=3/3) in levels comparable to control brains (Figure 9A, n=4/4, Supplemental Figure 5A, n=5/5). We next asked whether the removal of *Jnk1* from the *Dlx5/6* territory alone could cause thalamocortical axon misrouting. In *Dlx5/6-CIE;Jnk1fl/fl* mice (Figure 9D), where *Jnk1* is conditionally removed from the *Dlx5/6* territory but *Jnk2* and *Jnk3* remain intact, thalamocortical axons crossed the DTB and entered the cortex as normal (Figure 9D, n=3/3). Thus, the removal of *Jnk2* and *Jnk3* constitutively, or the conditional removal of only one JNK gene, *Jnk1*, from the *Dlx5/6* territory, are themselves insufficient to cause thalamocortical axon misrouting.

**Figure 9.**
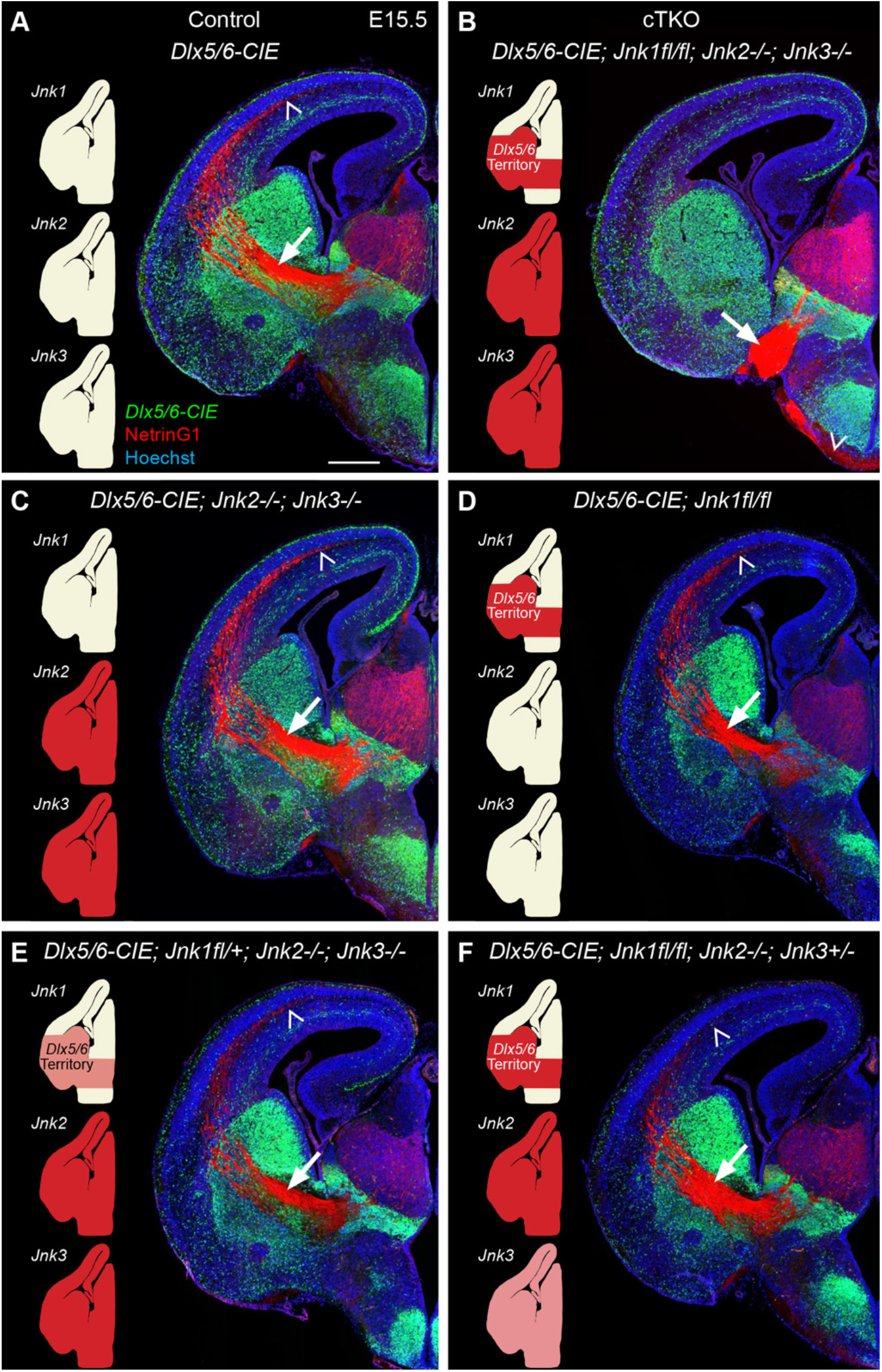
Thalamocortical axons require JNK signaling in the *Dlx5/6*-positive territory. A. NetrinG1-labeled thalamocortical axons project through the *Dlx5/6*-positive ventral forebrain in the internal capsule (arrow) in route to the cortex (open arrowhead). B. In *cTKO* brains, NetrinG1-positive axons project to the ventromedial margin of the telencephalon (arrow) and extend into the hypothalamus (open arrowhead). C-F. NetrinG1-postive thalamocortical axons project through the internal capsule (arrows) and into the cortex (open arrowheads) in *Dlx5/6-CIE;Jnk2-/-;Jnk3-/-* (C), *Dlx5/6-CIE;Jnk1fl/fl* (D), *Dlx5/6-CIE;Jnk1fl/+;Jnk2-/-;Jnk3-/-* (E), and *Dlx5/6-CIE;Jnk1fl/fl;Jnk2-/-;Jnk3+/-* (F) mice. In schematics, red indicates complete loss of a gene, while pink indicates one copy of the gene is deleted. Scale bar: 400 µm.

Since our data indicated that dosage of *JNK* gene function was an important component of the thalamocortical axon misrouting phenotype, and *JNK* genes are known to compensate for one another (Brecht et al., 2005; Chen et al., 2005; Jaeschke et al., 2006), we next asked whether retaining a single copy of either *Jnk1* or *Jnk3* was sufficient for proper thalamocortical axon extension. In *Dlx5/6-CIE;Jnk1fl/+;Jnk2-/-;Jnk3-/-*mice (Figure 9E), only one copy of *Jnk1* is conditionally removed from the *Dlx5/6* lineage in a *Jnk2/Jnk3* double knockout background. Thalamocortical axons were able to extend across the DTB and into the cortex of these mice at E15.5 (Figure 9E, n=5/5) and P0 (Supplemental Figure 5D, n=4/4), suggesting that only one copy of *Jnk1* is needed in the *Dlx5/6* territory for proper thalamocortical axon extension. Similarly, in *Dlx5/6-CIE;Jnk1fl/fl;Jnk2-/-;Jnk3+/-*mice, where only one copy of *Jnk3* is present, thalamocortical axons extended properly though the internal capsule and into the cortex (Figure 9F, n=5/5). Retaining one copy of *Jnk1* in the *Dlx5/6* territory, or one copy of *Jnk3* in the mouse, is sufficient for thalamocortical axons to cross the DTB and reach the cortex. Thus, it is the complete loss of JNK function from the *Dlx5/6* territory that leads to the non-autonomous misrouting of thalamocortical axons in *cTKO* mice.

## Discussion

In the current study, we demonstrate a novel, non-autonomous requirement for JNK signaling in thalamocortical axon pathfinding. In *cTKO* mice, *Jnk1* is conditionally removed from *Dlx5/6* lineage cells in a *Jnk2/Jnk3* null background, which completely eliminates JNK signaling from the territory through which thalamocortical axons traverse. Beginning early in development, thalamocortical axons are unable to cross the diencephalon-telencephalon boundary (DTB) in *cTKO* brains, and instead misroute ventrally into the hypothalamus and rostrally along the ventromedial margin of the telencephalon. Other longitudinal axon tracts in the *cTKO* brain, including the corticothalamic, striatonigral, and nigrostriatal axons, are also unable to cross the DTB, and fail to reach their intended targets through normal trajectories. The Islet1-positive corridor and Nkx2.1-positive globus pallidus cells are mispositioned in the *cTKO* ventral telencephalon, and both fail to extend caudally to the DTB region. Guidepost cells in the ventral telencephalon which normally extend axons to the dorsal thalamus are severely reduced in number in the *cTKO* brain. An allelic series of different mouse mutants reinforced the non-autonomous requirement for JNK signaling in thalamocortical axon pathfinding and demonstrated that all three *JNK* genes must be removed from the *Dlx5/6* territory for thalamocortical axon misrouting to occur.

### JNK signaling in the Dlx5/6 territory is essential for the establishment of major forebrain and midbrain axon tracts

Our data strongly suggests that correct thalamocortical axon pathfinding relies on intact JNK signaling within the *Dlx5/6* territory. The *Dlx5/6* expression domain spans much of the thalamocortical axon pathway, encompassing portions of the prethalamus and hypothalamus in the diencephalon, as well as the developing striatum, globus pallidus, and corridor region of the ventral telencephalon (Panganiban and Rubenstein, 2002; Stenman et al., 2003). Thus, the *Dlx5/6* territory contains many axon guidance cues and intermediate target cells that normally instruct thalamocortical axon guidance, and these cells or the factors they produce may be compromised by the conditional deletion of JNK in *cTKO* mice.

For instance, crossing of the DTB by thalamocortical axons is mediated in part by chemorepulsion from the hypothalamus (Anton-Bolanos et al., 2018). Hypothalamic-expressed Slits act as chemorepellents for thalamocortical axons to direct them away from the midline and steer them towards the ventral telencephalon (Bagri et al., 2002; Braisted et al., 2009; Lopez-Bendito et al., 2007). In *Slit2* and *Slit1/Slit2* knockout mice, thalamocortical axons misroute ventrally into the hypothalamus, and in *Slit1/Slit2* knockout mice, nigrostriatal axons also ectopically project into the hypothalamus (Bagri et al., 2002). Similarly, both thalamocortical and nigrostriatal pathways misproject ventrally to form U-shaped bundles beneath the hypothalamus in *cTKO* mice. This raises the possibility that loss of JNK signaling in the *Dlx5/6* domain of the hypothalamus may compromise Slit function in the developing *cTKO* brain. Although JNK has been shown to act upstream of Slit signaling in Drosophila (Iida et al., 2019; Vaughen and Igaki, 2016), connections between these pathways have not yet been identified in mice.

The ventral telencephalon also contains numerous intermediate target cells that facilitate thalamocortical axon pathfinding. Corridor cells are one population of *Dlx5/6* cells in the ventral telencephalon known to non-autonomously influence thalamocortical axon guidance. The telencephalic corridor is formed by a subset of Islet1-positive lateral ganglionic eminence cells that migrate ventromedially to form a permissive territory located between the Nkx2.1-positive medial ganglionic eminence and globus pallidus (Lopez-Bendito et al., 2006). The migration of corridor cells is aided in part by Slit2 expression in the ventral telencephalon, and when disrupted, corridor cells do not migrate caudally, and thalamocortical axons fail to cross the DTB (Bielle et al., 2011). In *cTKO* mice, both Islet1- and Nkx2.1-positive cells fail to extend caudally towards the DTB in the telencephalon, and Islet1-positive cells ectopically infiltrate the developing globus pallidus. Since disruptions to the corridor occur in other mouse mutants with thalamocortical axon misrouting phenotypes (Backer et al., 2018; Bielle et al., 2011; Lopez-Bendito et al., 2006; Morello et al., 2015; Simpson et al., 2009; Uemura et al., 2007), it is plausible that the mispositioning of corridor cells in *cTKO* mice leads to misrouting of thalamocortical axons.

In addition to corridor cells, early pioneering axons from neurons located in the ventral forebrain may assist the growth of thalamocortical axons. For instance, striatal axons have been proposed to serve as scaffolding for thalamocortical axons to cross the DTB, and indeed, in *OL-protocadherin-/-, Frizzled3-/-*, and *Celsr3-/-*mice, both striatonigral and thalamocortical axons fail to cross the DTB (Jia et al., 2014; Morello et al., 2015; Uemura et al., 2007). In *cTKO* mice, Darpp32-positive striatal axons fail to extend past the DTB, and instead coalesce in the ventral telencephalon. JNK is known to be important in neurite architecture (Coffey et al., 2000), neuronal polarity (Oliva et al., 2006), axon guidance (Shafer et al., 2011), and axon regeneration (Raivich et al., 2004), and therefore could play a role in the extension of striatal axons. Conditional removal of JNK function solely from the striatum would help determine if striatal axons have an autonomous requirement for JNK signaling, and if the striatal defects could be responsible for other axon misrouting phenotypes in *cTKO* mice.

Guidepost neurons, which extend axons from the ventral forebrain to the dorsal thalamus, are another population of ventral telencephalic cells thought to be important in early thalamocortical axon pathfinding (Braisted et al., 1999; Metin and Godement, 1996; Molnar and Cordery, 1999). In *cTKO* mice, guidepost neurons, identified by retrograde labeling from the thalamus, are severely reduced in number, similar to other mutants in which thalamocortical axons fail to extend across the DTB (Jones et al., 2002; Lakhina et al., 2007; Mitsogiannis et al., 2017; Tuttle et al., 1999; Uemura et al., 2007). Interestingly, guidepost neurons are *Dlx5/6*-positive, and when *Celsr3* was conditionally removed from the *Dlx5/6* territory, these cells were missing (Feng et al., 2016). Unraveling which populations of cells within the *Dlx5/6*-lineage require JNK function to facilitate thalamocortical axon pathfinding will be a major future effort.

### Non-canonical WNT signaling in thalamocortical axon guidance

The Planar Cell Polarity (PCP) pathway is a component of non-canonical Wnt signaling, and is known for its roles in tissue morphogenesis, cilia organization, and directed cell migration (Simons and Mlodzik, 2008; Wang and Nathans, 2007). Two seven-transmembrane receptors for Wnt ligands in the PCP pathway, *Frizzled3* (Goodrich and Strutt, 2011; Vinson et al., 1989) and *Celsr3* (Usui et al., 1999), play essential roles in axon guidance (Tissir et al., 2005; Wang et al., 2002), however the underlying mechanism is currently unknown. In both *Frizzled3* and *Celsr3* knockout mice, there are severe disruptions to major axon pathways including the thalamocortical, corticothalamic, and nigrostriatal tracts, and a near complete breakdown of the internal capsule (Fenstermaker et al., 2010; Tissir et al., 2005; Wang et al., 2002; Wang et al., 2006). These findings are very reminiscent of the *cTKO* mouse, in which thalamocortical and corticothalamic axons fail to extend properly through the telencephalon, and nigrostriatal axons are misrouted at the level of the hypothalamus. *Frizzled3* and *Celsr3* have also been conditionally removed using the *Dlx5/6* driver line, and nearly identical phenotypes to the constitutive deletions of both genes have been reported (Hua et al., 2014; Qu et al., 2014; Zhou et al., 2008; Zhou et al., 2009), suggesting it is the expression of PCP proteins in the *Dlx5/6* territory that is critical for proper axon guidance. Interestingly, JNK signaling is thought to be a major downstream effector of the WNT/PCP pathway (Niehrs, 2012). Thus, it is possible that PCP proteins act upstream of JNK in *Dlx5/6*-positive cells to non-autonomously regulate axon guidance in the developing forebrain.

Recent studies have looked more closely at the intermediate target territories in both the *Celsr3* and *Frizzled3* mutants. Ventral telencephalic guidepost cells were missing or severely reduced in *Celsr3* (Jia et al., 2014) and *Dlx5/6;Celsr3fl/fl* (Feng et al., 2016) mice, respectively, similar to the severe reduction observed in the *cTKO* forebrain. In *cTKO* mice, telencephalic Islet1-positive corridor cells invade the globus pallidus, which was also observed in *Celsr3* (Jia et al., 2014) and *Frizzled3* (Morello et al., 2015) mutants. *Iselt1*-positive cells in the prethalamus, another *Dlx5/6*-positive domain, fail to extend across the DTB in *cTKO* and *Islet1Cre;Celsr3fl/fl* mice (Feng et al., 2016). Additionally, in both the *Celsr3* and *cTKO* mutants, Nkx2.1-positive cells failed to extend caudally to the DTB region (Jia et al., 2014). Finally, striatal connections were compromised in both *Celsr3-/-* and *Frizzled3-/-* mice (Jia et al., 2014; Morello et al., 2015), similar to Darpp32-positive axons failing to extend past the DTB in *cTKO* brains. While mechanisms underlying the interplay between intermediate target cells and growing axons from major pathways in the developing forebrain remain to be fully dissected, it will be important to determine whether JNK acts downstream of non-canonical WNT signaling to coordinate this process.

### Conclusions

Thalamocortical axons extend a long distance to reach their cortical targets, and disruptions at any point along their trajectory can lead to lasting changes in neural connectivity and function. Here, we identified a novel, non-autonomous requirement for JNK signaling in the early pathfinding of thalamocortical axons. Elucidation of upstream and downstream regulators of JNK function in this critical developmental process will advance our understanding of thalamocortical connectivity, and in turn, inform us on how neurodevelopmental diseases may arise.

## Materials and Methods

### Animals

The Office of Laboratory Animal Resources at West Virginia University housed and cared for mice (*Mus musculus*) used in these studies. All mouse procedures were approved by and performed in accordance with the Institutional Animal Care and Use Committee at West Virginia University. Mice in all crosses were acquired and maintained on a C57BL/6J background. The individual mouse strains are as follows: C57BL/6J (Stock # 000664, The Jackson Laboratory); *Dlx5/6*-Cre-IRES-EGFP (*Dlx5/6-CIE*; (Stenman et al., 2003), floxed Mapk8tm1Rjd mice (*Jnk1fl/fl*; Das et al., 2007); kindly provided by Dr. Roger Davis); Mapk9tm1Flv (*Jnk2-/-*; Stock # 004321, The Jackson Laboratory); Mapk10tm1Flv (*Jnk3-/-*; Stock # 004322, The Jackson Laboratory). *Dlx5/6-CIE* control animals were generated by crossing C57BL/6J dams with *Dlx5/6-CIE* hemizygous males. *Jnk1fl/fl;Jnk2-/-;Jnk3-/-* dams were mated with *Dlx5/6-CIE*;*Jnk1fl/+;Jnk2-/-;Jnk3+/-*males to generate *Dlx5/6-CIE;Jnk1fl/fl;Jnk2-/-;Jnk3-/-*), *Dlx5/6-CIE;Jnk1fl/+;Jnk2-/-;Jnk3-/-*, and *Dlx5/6-CIE;Jnk1fl/fl;Jnk2-/-;Jnk3+/-* mice. *Jnk2+/-;Jnk3-/-* dams were mated with *Dlx5/6-CIE;Jnk2-/-;Jnk3+/-* males to generate *Dlx5/6-CIE;Jnk2-/-;Jnk3-/-* animals. *Jnk1f/fl* dams were mated with *Dlx5/6-CIE;Jnk1fl/+* males to generate *Dlx5/6-CIE;Jnk1fl/fl* animals.

### Tissue collection and processing

Dams were placed into male cages each night and checked every morning for a vaginal plug. The day of the vaginal plug was recorded as embryonic day 0.5. Timed-pregnant dams were euthanized, embryos were collected in 1X phosphate-buffered saline (PBS; 136.9mmol NaCl, 2.683mmol KCl, 4.290mmol Na2HP04 7H2O, 1.470mmol KH2P04), and tissue was taken for genotyping. E12.5 and E13.5 whole heads were submerged in 4% paraformaldehyde in PBS (PFA; Millipore Sigma, 818715). E15.5 and E17.5 brains were dissected, then submerged in 4% PFA. For P0 collections, newborn pups were deeply anesthetized and transcardially perfused with 1X PBS followed by 4% PFA. P0 brains were dissected and submerged in 4% PFA. All material was fixed for 24 hours in 4% PFA at 4°C before being passaged through a sucrose series (10%, 20%, 30%) for cryoprotection. Material was then embedded in Tissue Freezing Medium (VWR 15146-019), frozen, and stored at -80°C. Frozen material was sectioned at 12 µm on a Leica cryostat (CM3050S) onto superfrost plus slides (Fisher, 12-550-15), and stored at -20°C prior to staining.

### Immunohistochemistry and Imaging

Slides were outlined with a pap pen (Electron Microscopy Sciences, 71310), and rehydrated for 20 mins with 1X PBS, followed by blocking buffer containing permeabilization solution (Myers et al., 2014) with either 5% normal donkey (EMD Millipore, S30-100ml) or goat serum (Thermofisher scientific, 16210064). Primary antibody was diluted in block, placed on the slides, and incubated overnight at 4°C. Slides were then rinsed with 1X PBS, followed by incubation with secondary antibodies diluted in block for 2 hours at room temperature. After incubation, slides were rinsed again with 1X PBS and counterstained with Hoechst (Thermo Scientific 62249, 1µg/ml) for 10 minutes. Slides were rinsed before being cover-slipped with mounting media (Mowiol 4-88, Polysciences, 17951) containing an anti-fade reagent (p-Phenylenediamine, Fisher Scientific, AC130570050). All slides were stored at 4°C prior to imaging. Brain sections were imaged using an Olympus VS120 Slide Scanner with a UPLSAPO 10x objective, as well as on a Zeiss 710 confocal microscope with a 20x Plan-Apo objective lens.

Primary Antibodies:

| **Host:** | **Antigen:** | **Concentration:** | **Company:** | **Product Number:** |
| --- | --- | --- | --- | --- |
| Rabbit | Calbindin | 1:2000 | Swant | CB38 |
| Rabbit | Darpp-32 | 1:1000 | Synaptic Systems | 382 002 |
| Chicken | GFP | 1:1500 | Abcam | ab13970 |
| Goat | Islet1 | 1:500 | R&D Systems | AF1837 |
| Rabbit | L1 | 1:250 | Aviva Systems Biology | ARP63103 |
| Goat | NetrinG1 | 1:250 | R&D Systems | AF1166 |
| Rabbit | Nkx2.1 | 1:500 | Santa Cruz | sc-13040 |
| Rabbit | p-JNK | 1:1000 | Promega | V7931 |
| Mouse | Tag1 | 1:12.5 | DSHB | AB_2315433 |
| Chicken | TH | 1:1000 | Aves Labs | TYH |

Secondary Antibodies:

| Antibody: | Concentration: | Company: | Product Number: |
| --- | --- | --- | --- |
| Donkey anti-Chicken 488 | 1:4000 | Jackson Immuno Research | 703-545-155 |
| Goat anti-Chicken 488 | 1:4000 | Invitrogen | A11039 |
| Donkey anti-Goat 546 | 1:2000 | Invitrogen | A11056 |
| Goat anti-Chicken 546 | 1:2000 | Invitrogen | A11040 |
| Goat anti-Mouse 546 | 1:2000 | Invitrogen | A11003 |
| Goat anti-Rabbit 546 | 1:2000 | Invitrogen | A11010 |
| Donkey anti-Chicken 647 | 1:2000 | Jackson Immuno Research | 703-605-155 |
| Donkey anti-Rabbit 647 | 1:2000 | Jackson Immuno Research | 711-605-152 |
| Goat anti-Rabbit 647 | 1:2000 | Invitrogen | A21244 |

### DiI labeling

#### E13.5

Dissected heads from E13.5 embryos were fixed in 4% PFA overnight at 4°C. Following fixation, heads were rinsed into 1X PBS. Whole heads were hemi-sected along the mid-sagittal plane, and a DiI crystal (Invitrogen, D3911) was placed into the dorsal thalamus of both halves of the brain. DiI-labeled heads were incubated at 37°C for 2 weeks in the dark in 1X PBS. Following incubation, each hemi-sected head was embedded in 3% agarose (Fisher Scientific, BP165-25), sectioned in the coronal plane at 150 µm on a Leica vibratome (VT1000S), and slide mounted in series. Two slices from each brain were used for quantification of retrogradely-labeled cells in the ventral telencephalon. The section in which thalamocortical axons were at the diencephalon-telencephalon boundary region was selected as the caudal position, and the slice immediately rostral to this location was used as the rostral position. The total number of DiI-positive cells located within the ventral telencephalon of each slice were counted in both the rostral and caudal positions in n=6 brains per genotype, and overall statistical differences were determined by Student’s *t*-tests.

#### E15.5

E15.5 brains were fixed in 1.5% PFA overnight at 4°C. After rinsing into 1X PBS, brains were embedded in 3% agarose and then sectioned coronally at 300 µm on a vibratome. DiI crystals were placed into one half of the dorsal thalamus in each of the slices. DiI-labeled brain slices were incubated for 3 days at room temperature in 1X PBS in the dark before being mounted onto slides.

#### P0

P0 mice were perfused and brains were stored in 4% PFA for 1 week at 4°C. Brains were then rinsed into 1X PBS and hemi-sected along the mid-sagittal plane. DiI crystals were placed into the thalamus of the right hemisphere, and into the cortex of the left hemisphere. DiI-labeled brains were incubated at 37°C for 8 weeks in the dark. Each half of the brain was embedded in 3% agarose, sectioned at 200 µm on a vibratome, then slide mounted before imaging. DiI-labeled brains from all timepoints were imaged with a Zeiss 710 confocal microscope using a 20x Plan-Apo objective lens.

#### CT scans

P0 mice were perfused and heads were fixed in 4% PFA for 2 days at 4°C, after which they were transferred to stability buffer (4% w/v paraformaldehyde (pH 7.2), 4% w/v acrylamide (Bio-Rad 1610140), 0.05% w/v bis-acrylamide (Bio-Rad 1610142), 0.25% w/v VA044 initiator (Fisher Scientific, NC0632395), 0.05% w/v Saponin (EMD Millipore, 558255), in 1x PBS) for 3 days at 4°C (Hsu et al., 2016). The samples underwent nitrogen desiccation in stability buffer (3 minutes vacuum, 3 minutes Nitrogen, 3 minutes vacuum), and then were placed in a water bath at 37°C for 3 hours. Heads were carefully removed from the stability buffer, and placed in 0.1 N iodine (Thermo Fisher Scientific, AC12422) to stain for 2 days, with iodine being replaced at 24 hours. The heads were embedded in 3% agarose, then were imaged on a Bruker SkyScan 1272 MicroCT scanner (Al 0.5 mm filter, 850 ms exposure, and 7 μm resolution).

## Acknowledgements

We would like to thank Amanda Ammer and Karen Martin for their excellent microscopy assistance. Imaging was performed in the WVU Imaging Facilities, which have been supported by the WVU Cancer Institute, the WVU HSC Office of Research and Graduate Education, and NIH grants P20RR016440, P30GM103488, U54GM104942, P20GM103434, and P30GM103503. This work was partially supported by a transition grant from the Office of Research and Graduate Education at WVU Health Sciences Center (EST) and graduate stipend support from NIH grant T32 GM133369 (JDS).

## Author Contributions

Jessica Cunningham, Conceptualization, Methodology, Formal analysis, Investigation, Writing-original draft preparation, Visualization; James Scripter, Formal analysis, Investigation; Stephany Nti, Investigation; Eric Tucker, Conceptualization, Methodology, Supervision, Writing-review and editing, Funding acquisition.

---

**Supplemental Figure 1.**
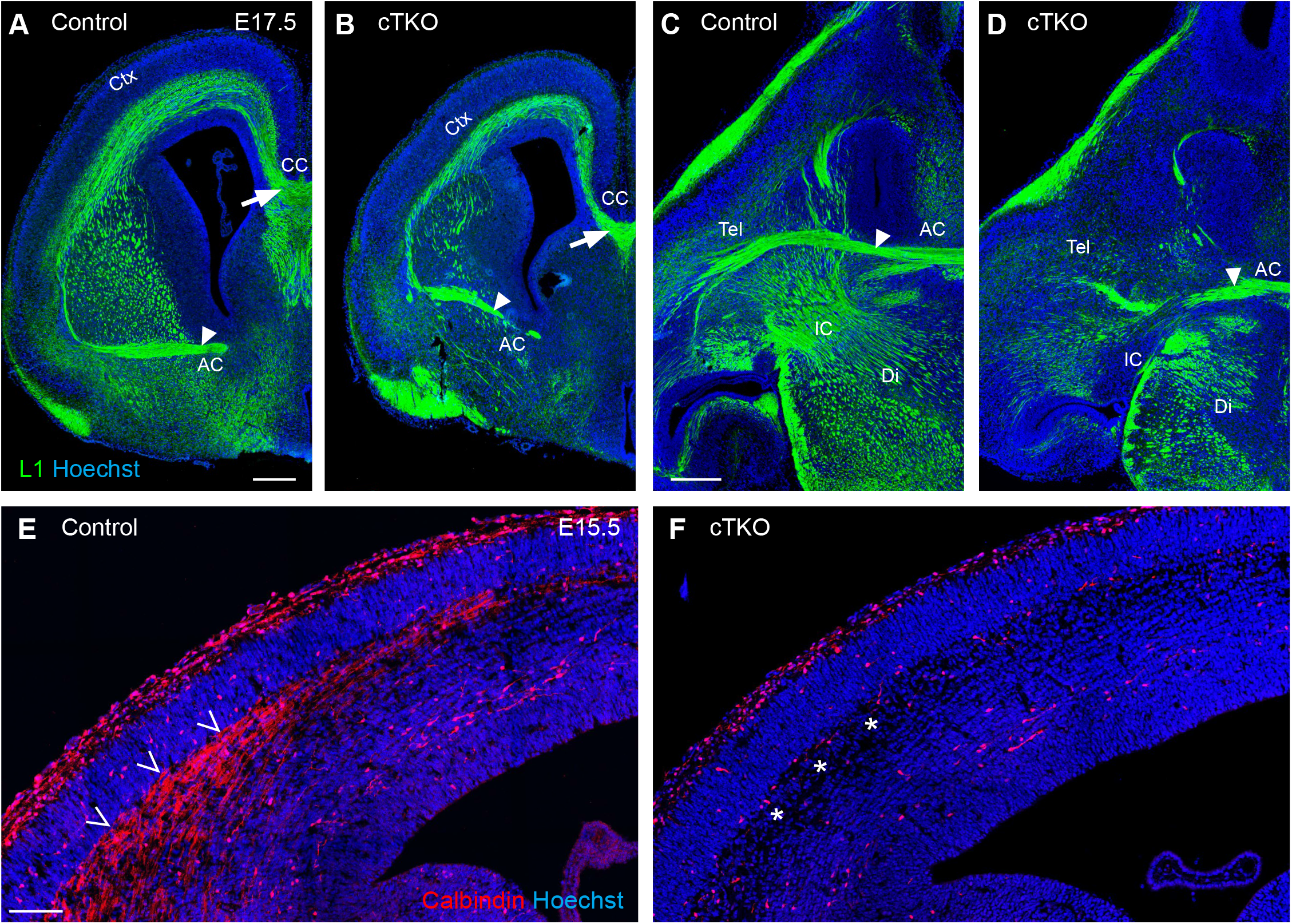
Commissural axon pathways are less disrupted than longitudinal pathways in *cTKO* forebrains. A-B. The corpus callosum (arrows) and anterior commissure (closed arrowheads) both cross the midline in control (A,C) and *cTKO* (B,D) brains. E-F. Immunohistochemistry for Calbindin in E15.5 brain sections labels a population of axons in the cortical wall of control brains (open arrowheads, E), which are missing from *cTKO* cortices (asterisks, F). Ctx- cortex, CC- corpus callosum, AC- anterior commissure, Tel- telencephalon, Di- diencephalon, IC- internal capsule. Scale bars: A-D 300 µm, E-F 100 µm.

**Supplemental Figure 2.**
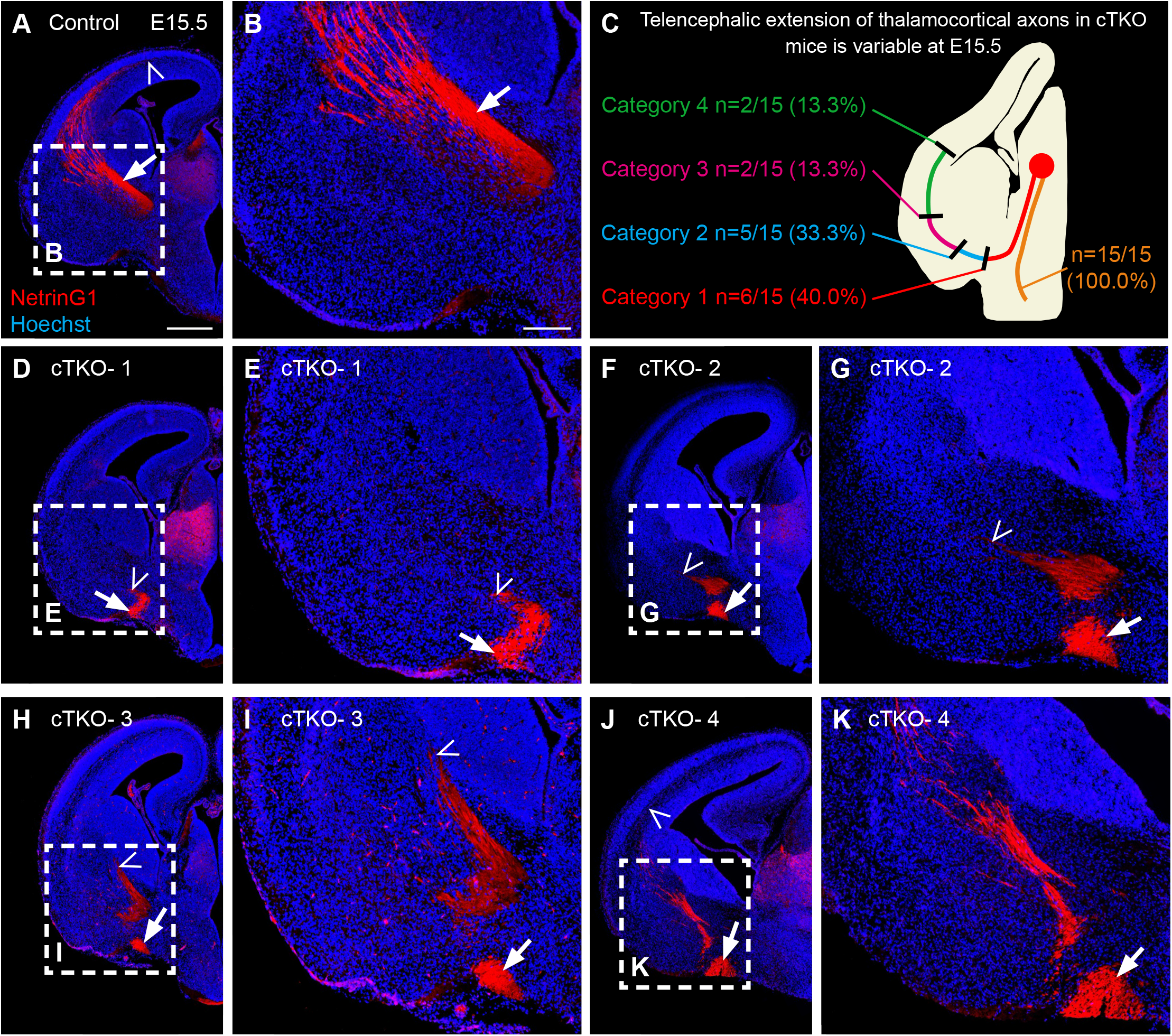
Thalamocortical axon misrouting is fully penetrant but the degree of ectopic entry into the telencephalon is variable in *cTKO* brains. A-B. In control brains, NetrinG1- positive thalamocortical axons project through the internal capsule (arrow) and into the cortex (open arrowhead). C-K. 15 *cTKO* brains were stained with NetrinG1 along the rostrocaudal axis at E15.5 to determine the penetrance of thalamocortical axon phenotyp es. C. Schematic summary of results. NetrinG1-positive axons aberrantly projected into the hyp othalamus in all 15 *cTKO* brains. Some *cTKO* brains contained NetrinG1 fibers that entered the telencephalon, but the degree that these fibers penetrated was variable (categories 1-4). D-E. In 6/15 *cTKO* brains, NetrinG1-positive axons (arrowheads) avoided entry into the ventral telencephalon, and instead coursed rostrally at the ventromedial margin of telencephalon (arrows; category 1). F-G. In 5/15 *cTKO* brains, most thalamocortical axons traveled at the ventromedial margin of the telecephalon (arrow), however some stray fibers (arrowheads) extended into the ventral telencephalon (category 2). H-I. In 2/15 *cTKO* brains, some axons (arrowheads) reached a rudimentary internal capsule but did not cross the pallial-subpallial boundary (category 3). J- K. In 2/15 *cTKO* brains, some axons (arrowheads) reached the cortical rudiment (category 4), however not to the same extent as controls. Scale bars: A,D,F,H,J 500 µm, B,E,G,I,K 200 µm.

**Supplemental Figure 3.**
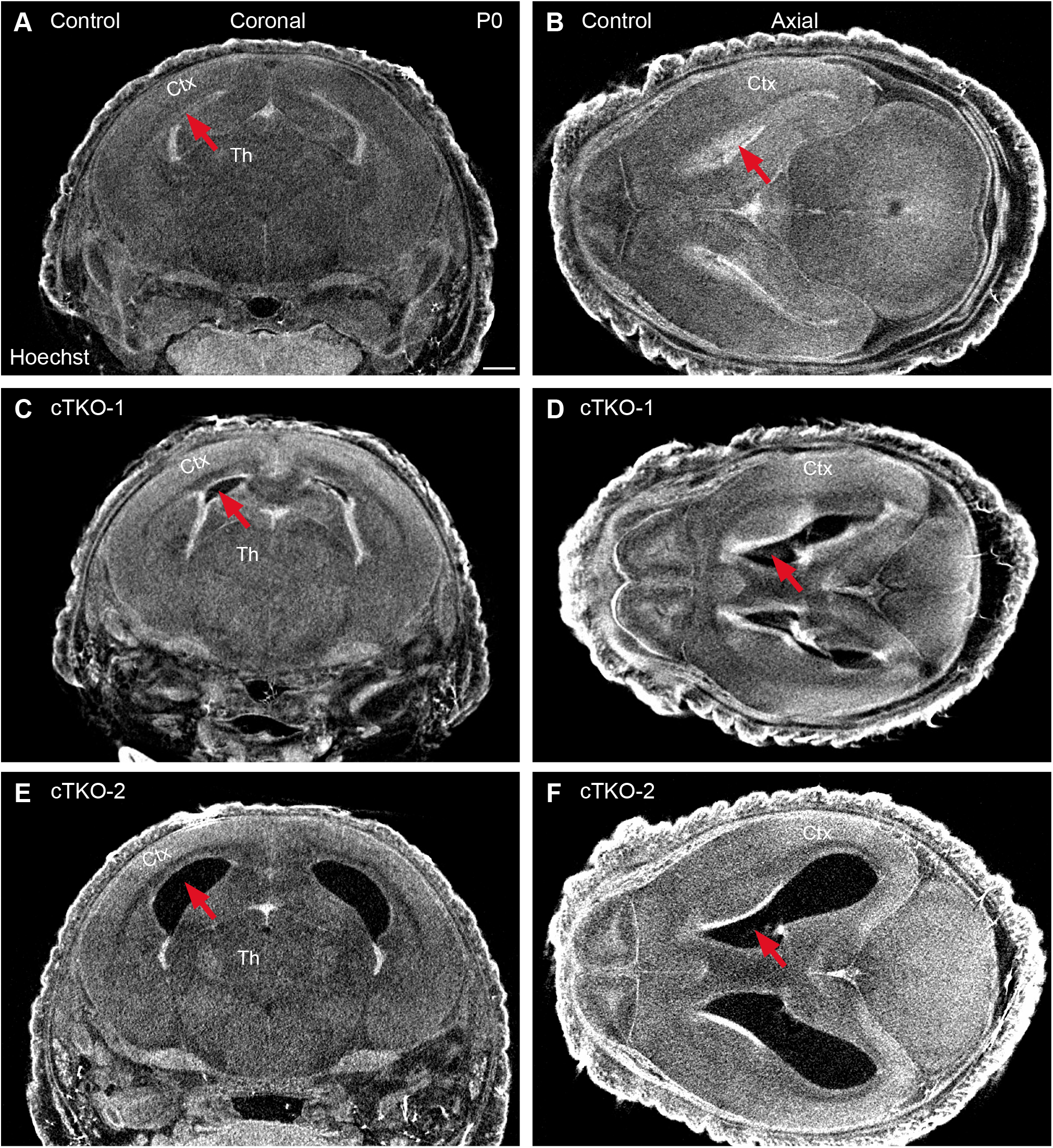
*cTKO* mice have enlarged ventricles at P0. A-B. In coronal (A) and axial (B) views of control brains, minimal ventricular space (arrows) is observed in CT images of whole heads at P0. C-F. Two different *cTKO* brains with enlarged lateral ventricles (arrows) evident in coronal (C,E) and axial (D,F) views. Ctx- cortex, Th- thalamus. Scale bar: 800 µm.

**Supplemental Figure 4.**
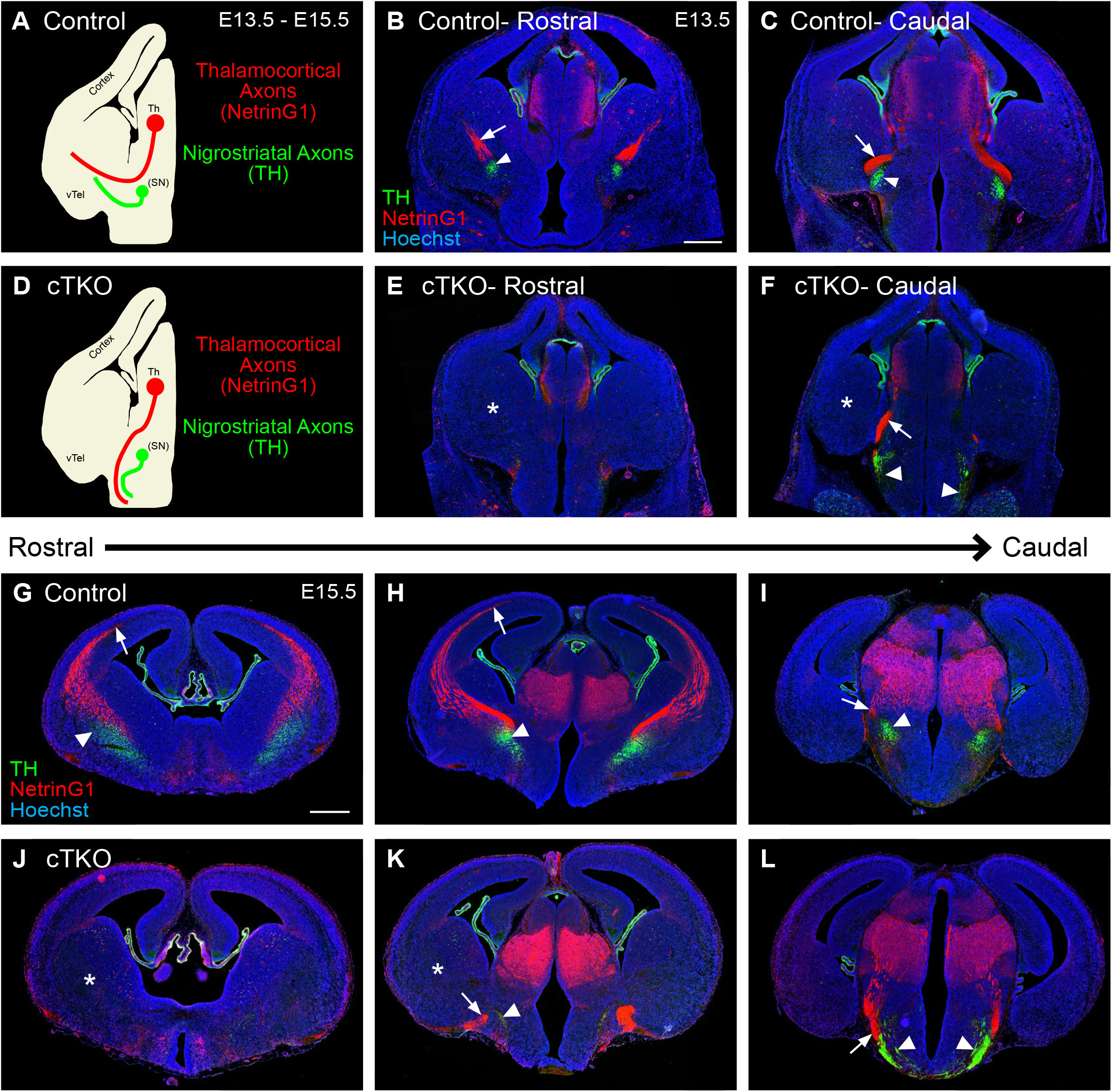
Nigrostriatal axons are misrouted in early development in *cTKO* mice. A-C. In control brains at E13.5, NetrinG1-positive (red) thalamocortical axons (arrows) and TH-positive (green) nigrostriatal (closed arrowheads) axons cross the DTB and enter the telencephalon. D-F. In E13.5 *cTKO* brains, both thalamocortical and nigrostriatal axons fail to reach the telencephalon (asterisks). Nigrostriatal axons are misrouted at E13.5 in *cTKO* brains (closed arrowheads). G-I. At E15.5, NetrinG1 axons (arrows) reach the cortex, and TH (closed arrowhead) axons are present in the striatum in control mice. J-L. In *cTKO* brains at E15.5, both sets of axons fail to reach the striatum (asterisk), and NetrinG1 (arrows) and TH (closed arrowheads) mis-project ventrally. Scale bars: B,C,E,F 400 µm, G-L 500 µm.

**Supplemental Figure 5.**
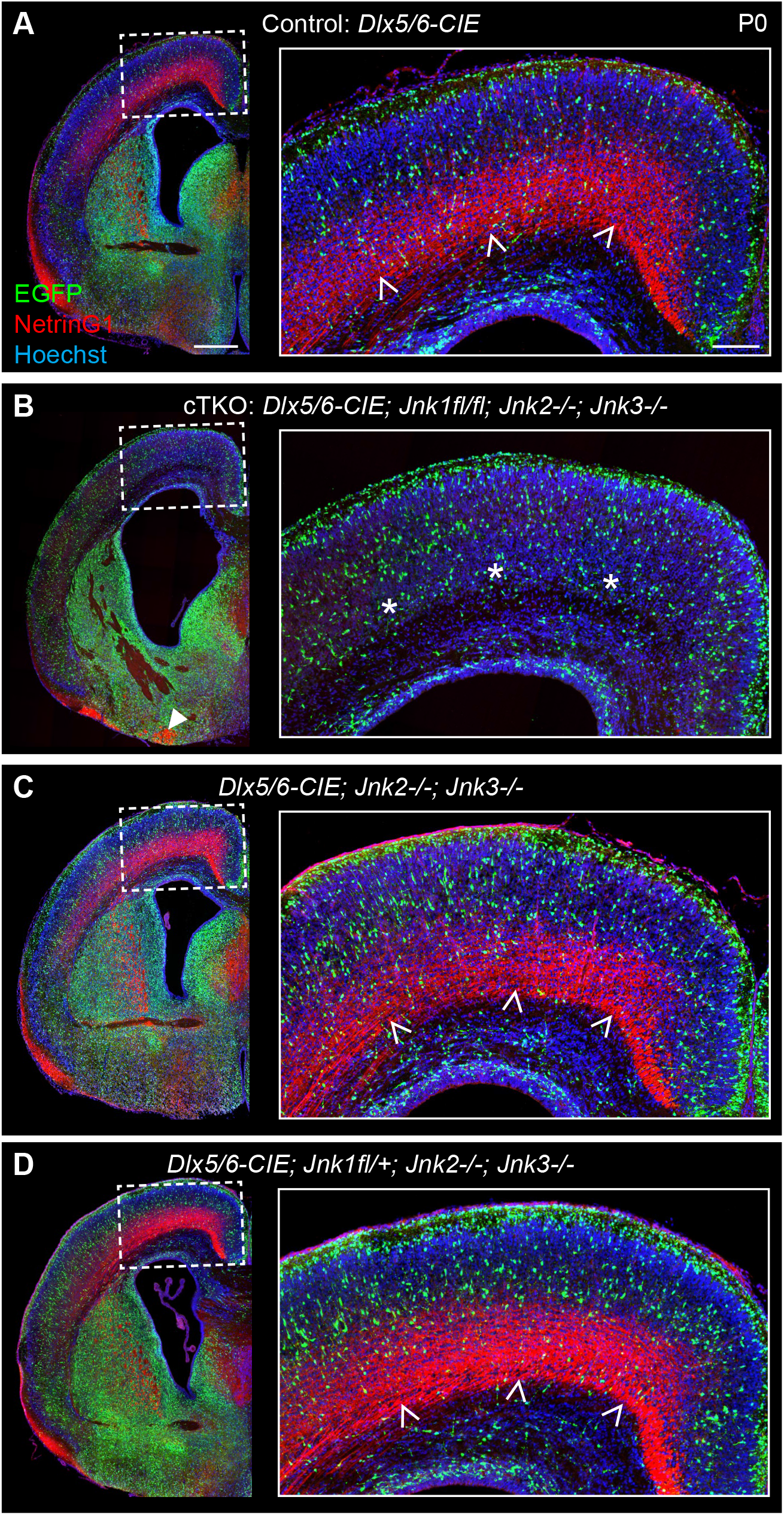
Thalamocortical axons are missing from the cortices of *cTKO* brains at P0. A. In control brains at P0, NetrinG1-positive thalamocortical axons are present in the cortical wall (open arrowheads). B. In *cTKO* brains, no NetrinG1-postive thalamocortical axons are present in the cortex (asterisks). An ectopic bundle of axons is present in the ventral telencephalon (closed arrowhead). C-D. In *Dlx5/6-CIE;Jnk2-/-;Jnk3-/-* (C) and *Dlx5/6-CIE;Jnk1fl/+;Jnk2-/-;Jnk3-/-* (D) mice, NetrinG1-positive thalamocortical axons are present in the cortical wall (open arrowheads), and no ectopic bundles are located in the telencephalon. Scale bars: 500 µm, insets 150 µm.

